# Circadian Disruption Elicits Sex-Specific Gut Microbiota, Endocannabinoidome and Lipid Mediator Responses

**DOI:** 10.64898/2025.12.22.695996

**Authors:** Pejman A. Pashaki, Timothy Niepokny, Tina Khalilzadehsabet, Elizabeth Dumais, Nicolas Flamand, Vincenzo Di Marzo, Eric M. Mintz, Cristoforo Silvestri

## Abstract

Circadian disruption is a pervasive environmental stressor that increases susceptibility to metabolic and inflammatory diseases, yet sex-specific adaptive strategies remain poorly understood. Here, we show that constant light (LL) exposure alters gut microbial communities and triggers sex- and tissue-specific host adaptations in the endocannabinoidome and other bioactive lipids. Using 16S rRNA sequencing, short chain fatty acid (SCFA) quantification, LC-MS/MS lipidomics, and cytokine profiling, we identified divergent coping strategies across brain, metabolic and intestinal tissues and reproductive organs. In females, LL induced microbial restructuring, with enrichment of *Rikenellaceae*, *Butyricicoccaceae*, and *Alistipes*, but these changes were uncoupled from short-chain fatty acids (SCFA) output. Also, they engaged *N*-acylethanolamine (NAE)-driven endocannabinoidome signaling in the brain (AEA, DHEA, OEA, PEA, SEA), accompanied by omega-6 prostaglandin upregulation and increased cytokines (IL-5, IFN-γ, MIP-2). The 2-monoacyl glycerols (2-MAGs) increased selectively in liver and skeletal muscle, reflecting tissue-specific lipid remodeling. In males, microbial shifts were limited (e.g., *Ruminococcaceae* depletion, *Tuzzerella* enrichment), yet LL triggered robust metabolic adaptation resulting in elevated SCFA levels (isobutyric, butyric, isovaleric, valeric acids) in faeces and elevation of several DHA-derived bioactive lipids in different intestinal tissues Few alterations in brain bioactive lipids were found, while several 2-MAGs were elevated skeletal muscles and testes. In contrast, several oxylipins were decreased within subcutaneous but not other adipose tissue depots. Together, the data shows that changes in bioactive lipid levels in response to circadian rhythm disruption are organ- and sex-specific as are alterations in microbiota populations, positioning sex as a key determinant of responses to circadian stress.

## Introduction

Environmental factors such as the photoperiod (duration of light exposure) significantly influence the behavior and metabolism of organisms. Light is the most significant factor regulating circadian rhythms, which are estimated to regulate between 20–50[% of the genome (1). Light detection by photosensitive retinal ganglion cells (ipRGCs) transfers information to the suprachiasmatic nucleus (SCN) in the hypothalamus to orchestrate circadian regulation (2). The majority of cells, including those in the SCN, contain a core clock mechanism involving genes such as CLOCK (Circadian Locomotor Output Cycles Kaput), BMAL1 (Brain and Muscle ARNT[like 1), CRY (Cryptochrome) and PER (Period). CLOCK and BMAL1 form a complex that drives PER and CRY gene expression, which in turn regulate their own expression, creating a cyclical loop (3). Disruption of the circadian rhythm by constant light or darkness in rodents induces weight gain, impaired glucose and triglyceride metabolism, reduced insulin sensitivity, altered energy expenditure (4, 5). Similarly, rotating light schedules that mimic shift work increase adiposity, hepatic lipid storage, glucose intolerance, and insulin resistance in mice (6). In humans, circadian disruption from shift work is linked to metabolic dysfunction and observed in night[shift workers, including nurses in Brazil and Poland, steel workers in Japan (7), as well as offshore oil industry workers in the North Sea (8). Even dim light at night can perturb metabolism by altering the timing of food intake (5) and energy expenditure (9), thereby increasing susceptibility to obesity (10), type 2 diabetes (11), and metabolic dysfunction-associated steatotic liver disease (MASLD) (12).

In obese individuals, chronic low-grade inflammation perturbs circadian clock gene expression (13) indirectly through hyperinsulinemia and hyperglycemia (14), and directly through cytokines (15) such as IL-6, IL-1β, TNF-α, and prostaglandins (including PGF_2α_ and PGE_2_), which can directly regulate BMAL1 expression in peripheral tissues (16, 17). The endocannabinoid system (ECS) regulates circadian rhythms via CB_1_/CB_2_ signaling, linking environmental cues to internal homeostasis (18, 19). Endogenous ligands of cannabinoid CB_1_ and CB_2_ receptors, known as endocannabinoids, show daily oscillations that align with sleep-wake cycles and may influence eating behavior (20). CB_1_ activation by synthetic cannabinoids disrupts circadian entrainment by light, while CB_1_ antagonism under conditions of constant dark results in subtle phase shifting (21). Evidence suggests that this may be a result of CB_1_ -mediated alteration of PER2 rhythm in the SCN (22) through altered levels of the endocannabinoid, 2-arachidonoyl-glycerol (2-AG) (19). The hypothalamic levels of this lipid are enhanced by blue light in mice to modulate orexin-A neuron firing in the perifornical hypothalamic area receiving inputs from the retinohypothalamic tract (23).

Beyond the ECS, the endocannabinoidome (eCBome) encompasses a broader family of lipid mediators structurally related to endocannabinoids, such as *N*-acylethanolamines (NAEs) and 2-monoacylglycerols (2[MAGs) (24). The eCBome displays circadian rhythmicity in the brain (25), jejunum (19), plasma, and liver (26), indicating broad regulation of the eCBome across tissues. For example, in the cerebellum, NAEs increase at light/dark transitions while 2-MAGs decrease, whereas in the cortex and hippocampus sleep modulates *N*-acyl phosphatidylethanolamine-specific phospholipase D (NAPE-PLD) expression and light exposure alters monoacylglycerol lipase (MAGL) (27). NAPE-PLD and MAGL, respectively, contribute to the formation and degradation of NAEs and 2-MAGs (28). Their levels also shift under stressful conditions (29), including chronic unpredictable mild stress (30) kainic acid induced excitotoxicity (31, 32), ischemia (33), inflammatory bowel diseases (IBD) (34), UV-B radiation exposure (35) and in obesity/diabetes (36). The eCBome, acts through G[protein[coupled receptors (GPCRs), establishing versatile signaling pathways (37). For example, *N*-palmitoyl-ethanolamine (PEA) activates GPR55 and peroxisome proliferator-activated receptor alpha (PPARα) (38); *N*-oleoyl-ethanolamine (OEA) activates TRPV1, PPARα and GPR119 (39), and *N*-docosahexaenoyl-ethanolamine (DHEA), whose levels are increased by dietary docosahexaenoic acid (DHA), acts through GPR110 and reduces cytokine production (40). DHA also mitigates HFD-induced circadian disruption, potentially through DHEA anti-inflammatory effects (41). Under prolonged metabolic (42), obesogenic, oxidative or inflammatory stress eCBome mediator production may be blunted (43, 44) or increased (45), and either imbalance disrupts homeostasis and elevates disease risk. Therefore, disruption of the eCBome under conditions of light cycle disruption may in part contribute to increased susceptibility to metabolic dysregulation.

Variation in most eCBome lipids and receptors have been associated with the gut microbiota, both dependent and independent of dietary lipid intake (46). The microbiota links diet to systemic physiology; for instance, diets rich in unsaturated fatty acids increase fecal levels of the corresponding eCBome mediators (47), and microbiota can convert eicosapentaenoic acid (EPA) and DHA into oxylipins or stearidonic acid (SDA) into its corresponding NAE (48). In this way, the microbiota translates dietary fat intake into metabolic responses partly through the production of eCBome lipids. Beyond diet, the microbiome also modulates host eCBome directly at both lipid and receptor expression levels[(49). The microbiota itself shows diurnal rhythmicity and influences circadian regulation in mice and humans; for example, *Lactobacillus reuteri* and *Dehalobacterium*, exhibit robust rhythmicity over time, while *Ruminococcaceae* (50) lose rhythmicity under jet-lag conditions, demonstrating the dynamic relationship between microbial activity and the host’s circadian clock. Microbial metabolites such as acetate, propionate and butyrate further affect peripheral clocks by modulating the expression of core clock genes including BMAL1 and PER2 in the peripheral tissues (51, 52).

Disruption of circadian rhythms may alter eCBome lipid levels and receptor-mediated signaling in a tissue specific manner, contributing to pathophysiological outcomes, such as increased susceptibility to metabolic disorders. To address the complex interplay between the eCBome, gut microbiota, and circadian rhythms, we investigated how constant light exposure in male and female mice affects the gut microbiome in feces and intestinal sections, as well as tissue-specific eCBome lipid levels at a defined circadian time point.

### Study design and methodology

Sixteen C57BL/6J mice (8 male/ 8 female), aged 33-weeks, were randomly assigned to one of two experimental groups (n=4 per group per sex) and housed individually with a running-wheel. Animals had ad libitum access to food (Prolab RMH 3000, 5P00; LabDiet, St. Louis, MO, USA) and water Animals were placed on a 12-hour light/12-hour dark (LD) cycle for 2-weeks before the beginning of this experiment. Half of the mice were maintained under the standard (12:12 - LD) photoperiod (control group) and half were exposed to constant light (24:0 - LL) (experimental group) for 10 consecutive days to challenge the circadian system with an extended photoperiod rather than to induce circadian rhythm disruption. Overhead lighting was approximately 100 lux, ambient chamber temperature was 21°±2°C, and wheel-running activity was monitored with Clocklab (Actimetrics circadian cabinets, Wilmette, IL, USA) to determine individual circadian rhythms. After 10 days in either LD or LL conditions, mice were transferred to clean, empty cages for feces collection at zeitgeber/circadian time (ZT/CT) 15-16. Feces were stored on dry ice during collection and at -80°C until ready for analysis. Following feces collection, mice were placed back in their respective lighting conditions for 5 days before being sacrificed between ZT/CT 11 by cervical dislocation.

Food was removed 6 hours prior to euthanasia and tissue collection was completed within 15-18 minutes. Animals were sacrificed over multiple successive days to account for time. Tissue dissection proceeded as follows: trunk blood collected in EDTA coated tubes and spun down for plasma collection, brain separated into the hypothalamus, cerebellum, left hemisphere, and right hemisphere (consisting of the cerebrum), intrascapular brown adipose tissue, liver (right lobe), epididymal/ovarian white adipose tissue, retroperitoneal white adipose tissue, inguinal subcutaneous white adipose tissue, gonads, soleus, gastrocnemius, quadriceps, colon (distal section with lumen content removed), caecum, ileum, jejunum, and duodenum. Tissues were weighed and 10-20mg/tissue was frozen in liquid nitrogen, moved to -80°C. The Institutional Animal Care and Use Committee (IACUC) at Kent State University approved all experiments and procedures (protocol number 488 EM 19-12) and they followed the National Institutes of Health guidelines for the care and use of laboratory animals for experimental procedures.

### Gut microbiota sequencing and library preparation

DNA Extraction and 16S rRNA Sequencing; total DNA was extracted using PowerSoil pro DNA Extraction kit (Qiagen, Hilden, Germany), following the manufacturer’s instructions. This included mechanical lysis steps by bead-beating and enzymatic digestion to ensure effective disruption of bacterial cell walls and maximize DNA yield. DNA quality and concentration were assessed using a plat reader and fluorometrically with a Qubit fluorometer (Thermo Fisher Scientific, MA, USA). A sequencing library was then prepared targeting the hypervariable V3–V4 regions of the 16S rRNA according to Illumina’s amplicon library preparation protocol for the MiSeq System using primer pair-F(5′-TCGTCGGCAGCGTCAGATGTGTATAAGAGACAGCCTACGGGNGGCWGCAG-3′),R(5′GTCTCGTGGGCTCGGAGATGTGTATAAGAGACAGGACTACHVGGGTATCTAATCC-3′), in combination with the Illumina DNA Prep kit (Sets A–D; Illumina, USA) and Illumina DNA Preparation kit (Illumina, USA) PCR amplification was performed under standardized cycling conditions, and amplicon products were purified using magnetic bead-based cleanup by AMPure XP beads. Indexed adapters were ligated during a second round of PCR to allow multiplexing of samples. Libraries were normalized and pooled at 4 nM, denatured, and diluted to a final concentration of 8 pM. Paired-end sequencing (2 × 300 bp) was performed using the MiSeq Reagent Kit v3 (600 cycles) on an Illumina MiSeq platform. Raw sequencing reads were processed using the DADA2 pipeline (53) which includes filtering for quality, dereplication, chimera removal, and inference of exact amplicon sequence variants (ASVs). Taxonomic classification was assigned on a reference database SILVA (54) corresponding to the targeted V3–V4 region. Data were processed using the MicrobiomeAnalyst platform (55). Microbiota composition was assessed by calculating alpha and beta-diversity indexes obtained using the Bray–Curtis index. Principal Coordinate Analysis (PCoA) was performed and analysed using the pairwise PERMANOVA analysis.

### Short-Chain Fatty Acid (SCFA) Analysis

SCFAs in feces were analyzed using gas chromatography with flame ionization detection (GC-FID). Briefly, 50–100 mg of feces was homogenized with 1 mL of Milli-Q water per 100 mg of feces, homogenized for 2 minutes, and centrifuged at 18,000 × g for 10 minutes at 4°C. Subsequently, 200 µL of the supernatant was transferred to a 2 mL Eppendorf tube, and 10 µL of the internal standard was added along with 200 µL of MTBE. The mixture was vortexed for 20 minutes and centrifuged at 18,000 × g for 10 minutes. Finally, 150 µL of the organic phase was used for GC-FID analysis.

### Lipid Extraction Protocol

For tissue samples, lipid extraction was performed using a modified Bligh and Dyer method (56). Briefly, weighed tissue samples were homogenized in an Eppendorf tube with 500 µl of Tris-HCl buffer (50 mM, pH 7) using a tissue homogenizer. The homogenate was transferred to a 15 ml glass tube, followed by the addition of 500 µl LC-MS grade methanol containing 0.01% acetic acid and 5 µl of the deuterated internal standards (ISTDs). One milliliter of chloroform was added with a glass pipette, and the mixture was vortexed for 30 seconds. Samples were centrifuged at 4,000 × g for 5 minutes at room temperature, and the organic phase (lower layer) was collected in a glass tube. This extraction step was repeated twice with 1 ml chloroform. The combined organic phases (∼3 ml chloroform) were evaporated to dryness using a rotary evaporator. Plasma (40 µl) samples we extracted as before(57). The dried lipid extracts of tissue and plasma samples were resuspended in 60 µl LC-MS mobile phase (50:50 A: B) and vortexed for 15 seconds and analyzed on a Shimadzu 8050 triple quadrupole mass spectrometer (58). Retention times of each compound was validated with a mix pure standard that was injected with each sample batch. For the MAGs, we combined the signal of *sn-*1(3) and *sn*-2 isomers, due to acyl migration from the *sn*-2 to the *sn*-1(3) position in presence of water. Only peaks with a signal-to-noise ratio ≥ 5 were kept for quantification, as recommended recently(59).

### Cytokines analyses

Cytokine and chemokine levels were determined via a high-sensitivity 18-plex discovery assay (MDHSTC18) performed by Eve Technologies (Calgary, AB).

### Statistical analysis

For gut microbiome analysis, data were processed using MicrobiomeAnalyst.ca (55). For lipid mediator (LC-MS/MS), SCFA, and cytokine datasets, statistical analyses were performed in Python (v3.11) with the scipy.stats package. Raw intensity values were log2-transformed to stabilize variance. Group comparisons were assessed using two-tailed independent-samples t-tests. Molecules were considered significantly regulated when p < 0.05 and the absolute log2 fold change exceeded 0.5.

## Results

### Constant Light Alters Overall Gut Microbial Community Structure

To investigate the impact of circadian rhythm disruption on gut microbiota, 16S rRNA gene sequencing was performed on fecal samples collected from male and female mice exposed to either a 12:12 light/dark cycle (LD) or constant light (LL). Stacked bar plots of genus-level relative abundances suggested shifts in the microbial community structure across experimental groups in a sex dependent manner. In females, LL exposure appeared to increase the proportion of *Lactobacillus*, *Eisenbergiella*, and *Clostridium_sensu_stricto_1*, accompanied by a decrease in *Turicibacter*. In males, LL exposure appeared to similarly induce a relative increase in *Lactobacillus* while decreasing *Clostridium_sensu_stricto_1* and *Lachnospiraceae_NK4A136* (Fig. 1A). Shannon α-diversity was not statistically significant between treatment groups in females; however, in males, a strong trend for decrease (P= 0.07) in α-diversity was found, which was maintained when data from both sexes were merged (Fig. 1 B). Analysis of β-diversity using Principal Coordinate Analysis (PCoA) based on Jensen-Shannon Divergence revealed that LL treatment significantly influenced microbial community composition in females (p = 0.03), but not in males (p = 0.43). In the merged analysis, LL also significantly shaped β-diversity (p = 0.035) (Fig 1C).

**Figure 1.**
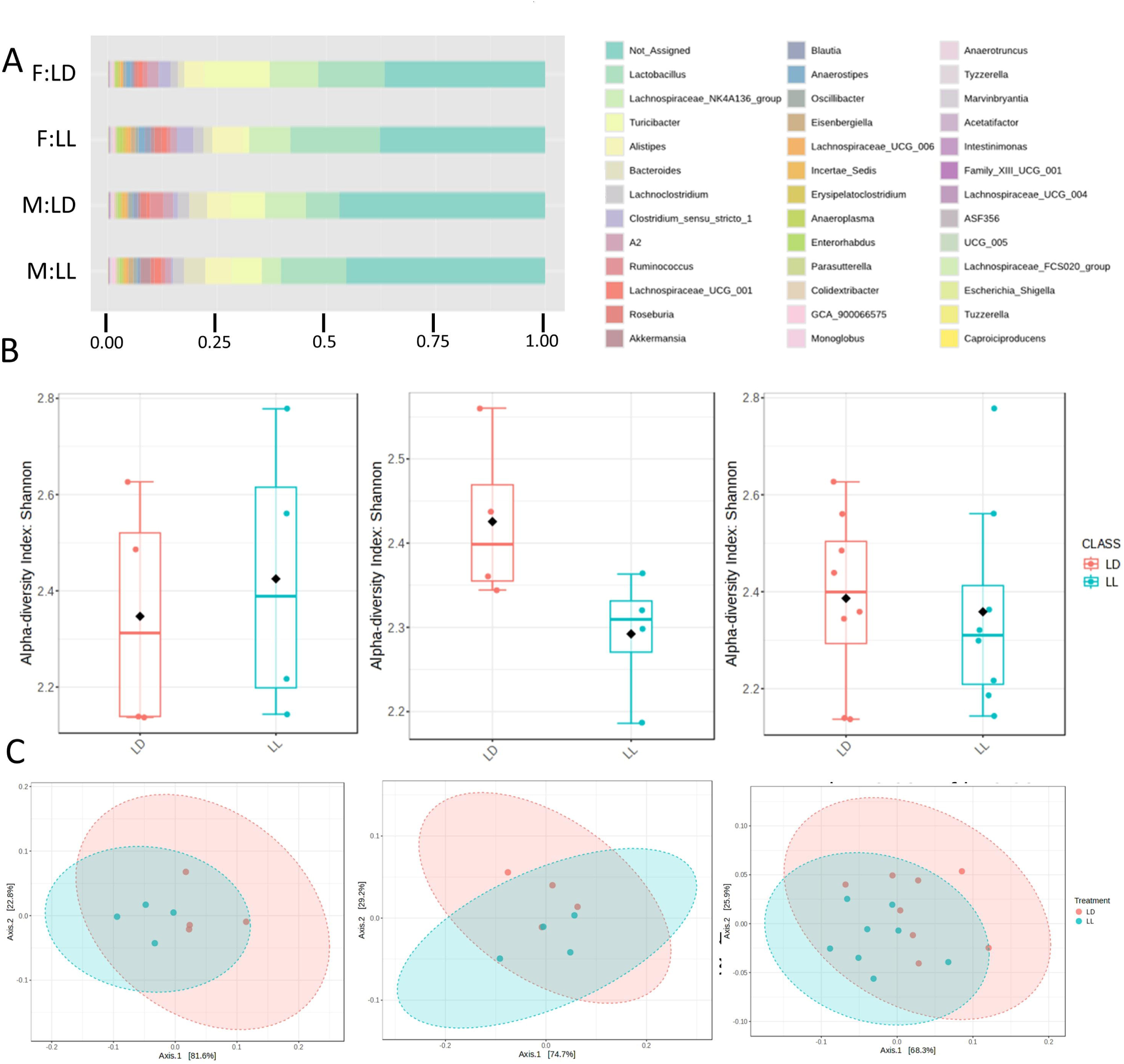
Effects of constant light on the gut microbiomes of male and female mice. (A) Stacked bar plots show relative abundance of the taxonomic composition at the genus level for female (F) and male (M) mice in light/dark (LD) constant light (LL). (B) Alpha diversity (Shannon index) of the gut microbiome at the genus level in females (left) p-value: 0.7, males (middle) p-value: 0.07, and combined samples (right) p-value: 0.7. (C) Beta diversity (PCoA using Jensen–Shannon divergence) showing microbial community differences in females (left) p-value: 0.04, males (middle) p-value: 0.4, and combined samples (right) p-value: 0.03.

Heat tree analysis (60) in males identified *Tuzzerella* as the only taxa that increased along with *Eisenbergiella*, whereas *Turicibacter* showed decreased. in females. Merged sex data revealed increases in *Eisenbergiella*, *Tuzzerella*, *Alistipes* (genus), and *Rikenellaceae* (family), and a significant reduction in *Ruminococcaceae UCG-005* (Fig. 2A). We then went on to perform LEfSe analysis, which further identified taxa that define the different treatments, and confirmed treatment-specific differences. In females, *Turicibacter* and *Alistipes*, were the abundant taxa that significantly changed in response to light treatment, with the former being high in LD and the latter high in LL mice, while males showed no significant taxa shifts. In merged data, *Turicibacter* and *Alistipes* were similarly identified along with decrease in *Lactobacillus* (Fig. 2B). We then identified effects of constant light exposure on the relative proportions of individual bacterial taxa (Fig. 2C). At the family level, females exhibited higher *Rikenellaceae* and *Butyricicoccaceae* with a marked reduction of *Erysipelotrichaceae*, while males displayed depletion of *Ruminococcaceae* under LL. In the merged sex analysis, *Rikenellaceae* remained enriched, while *Erysipelotrichaceae* was consistently depleted under LL. At the genus level, males showed increased *Tuzzerella* and *Oscillospirales-UCG-010* under LL, while females showed an increase in *Alistipes*, *Eisenbergiella*, *Enterorhabdus*, and *Butyricicoccus*, and *Turicibacter* and *Ruminococcaceae-UCG-005* decreased under LL. Merged sexes showed that *Alistipes*, *Eisenbergiella*, *Enterorhabdus*, and *Tuzzerella* were enriched, while *Turicibacter* and *Ruminococcaceae-UCG-005* were reduced, under LL.

**Figure 2.**
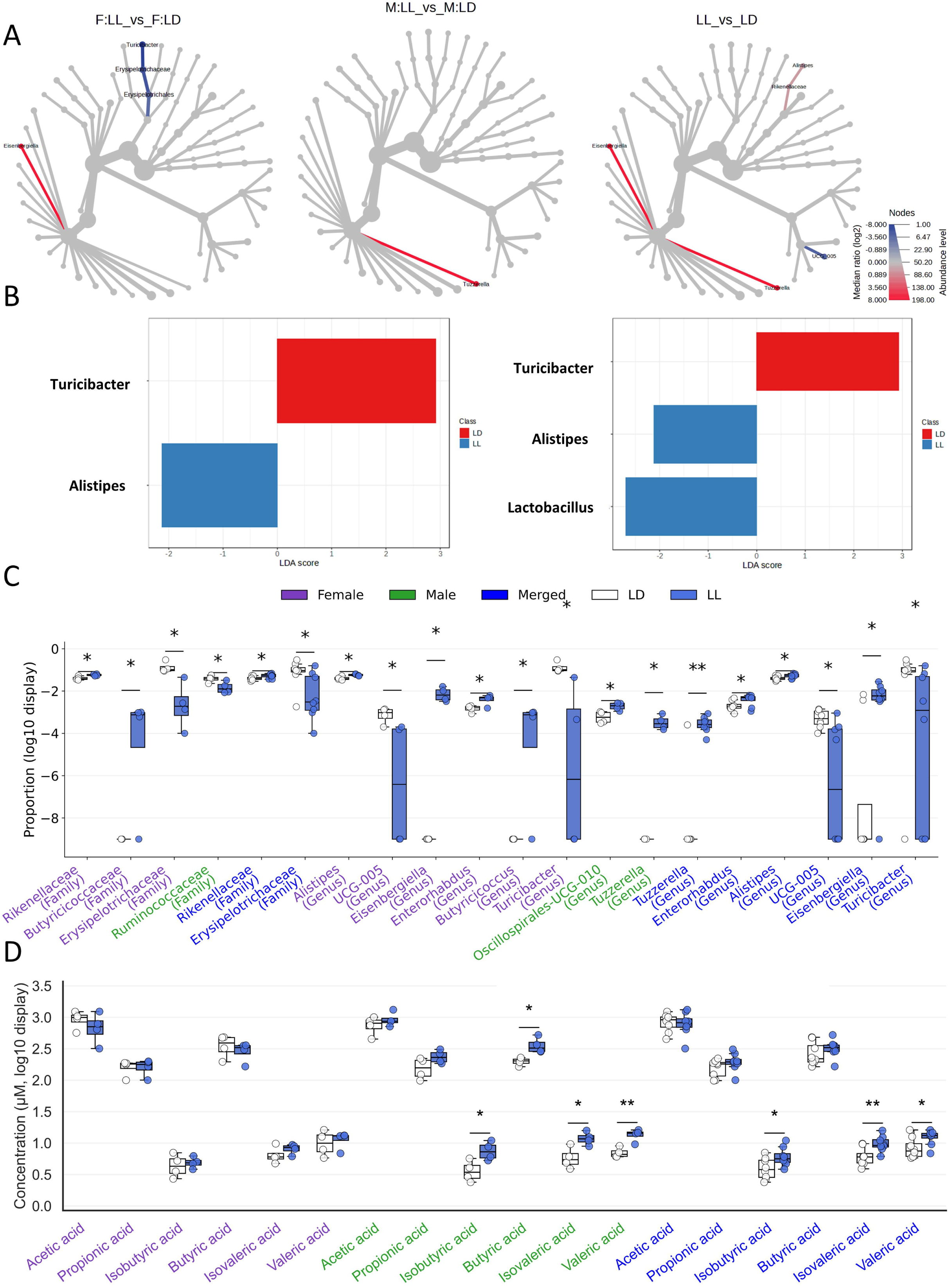
Differential abundance of microbial taxa between constant light (LL) and light/dark (LD) conditions at the genus level. A, Heat tree visualization shows taxa that significantly differ in abundance females (F; left), males (M; middle), combined sexes (right) between LL and LD groups. Red branches indicate taxa enriched under LL conditions, while blue branches indicate those more abundant in LD. Node size corresponds to taxon abundance, and color intensity reflects the log□-fold change in median relative abundance. B, Linear Discriminant Analysis Effect Size (LEfSe) analysis (P-value cutoff < 0.05, log LDA score > 2.0, the LDA score on the x-axis indicates the effect size of each taxon); C, Box-and-whisker plots of relative abundance for gut microbiota taxa (Family and Genus) that differed significantly between light–dark (LD) and constant light (LL) conditions; females (F; left; purple), males (M; middle; green), combined sexes (right; blue). Significant differences were determined by students t-test (p < 0.05). D, Box-and-whisker plots of Log10 fold change in the concentrations of various SCFAs that were measured via gas chromatography. The analysis was performed using an unpaired Student’s t-test separately for females (F; left; purple), males (M; middle; green), combined sexes (right; blue). Asterisks denote statistically significant differences: * p < 0.05.

Short chain fatty acids (SCFAs) were then measured via gas chromatography to determine if alterations in the gut microbiomes by LL modified the levels of these key bacterial metabolites in feces. Surprisingly, despite notable taxonomic shifts, LL did not in induce significant changes in SCFA levels in female mice; however, it did significantly increase isobutyric, butyric, isovaleric, and valeric acid in males. Except for butyric acid, these increases were similarly observed when data from male and female samples were merged. The results suggest a sex-dependent metabolic output, with males exhibiting greater fermentation activity under circadian disruption (Fig. 2D).

### Endocannabinoidome Lipid Mediators Are Altered by Constant Light

Principal Component Analysis (PCA) was used to assess global variation, with treatment separation tested by PERMANOVA (Euclidean distances; permutations with F, p, and R² reported), and variable contributions quantified by Cohen’s d per molecule. We first sought to investigate the global impact of LL exposure on the eCBome by PCA within anatomical groups. This analysis revealed significant treatment separation within the gastrointestinal tract (p=0.008) (liver, duodenum, jejunum, ileum, colon), brain regions (p=0.036) (cerebellum, hypothalamus, midbrain, right hemisphere), and skeletal muscle (p=0.001) (gastrocnemius, quadriceps, soleus). In contrast, adipose tissues (p=0.408) (gonadal WAT, retroperitoneal WAT, inguinal WAT, interscapular BAT) did not show significant differences. Also, PCA showed a significant sex effect on lipid profiles (Fig. 3), motivating subsequent analyses stratified by females, males, and merged data. This approach allowed us to capture the distinct remodeling of the eCBome within tissues in response to constant light.

**Figure 3.**
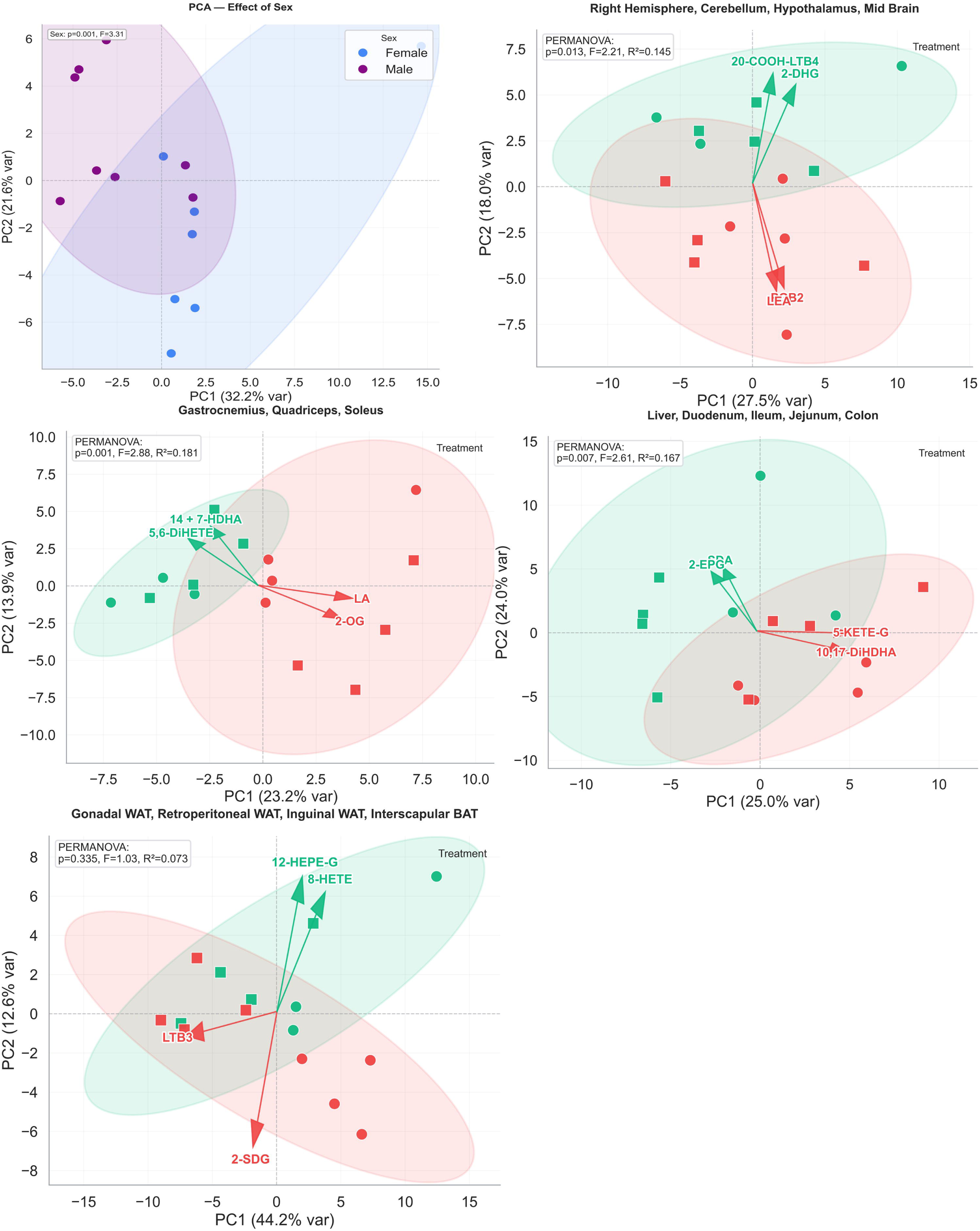
Principal Component Analysis (PCA) plots illustrate treatment- and sex-dependent clustering of lipid mediators in (A) Female (blue) and male (purple), there is significant sex effect in the groups (p= 0.001). PCA showed significant treatment effect in (B) gastrointestinal tract (p=0.008) (liver, duodenum, ileum, jejunum, colon), (D) skeletal muscle (p=0.001) (gastrocnemius, quadriceps, soleus), and (E) brain regions (p=0.036) (cerebellum, hypothalamus, midbrain, right hemisphere) but not significant in (C) adipose Tissues (p=0.408) (gonadal, retroperitoneal, inguinal, interscapular BAT). Ellipses represent 95% confidence intervals. PERMANOVA results (F, p, R²) indicate significant effects of treatment or sex. Arrows highlight the top two lipid mediators contributing to separation along PC1 and PC2.

Constant light (LL) exposure induced extensive remodeling across all major classes of lipid mediators. To highlight coordinated physiological patterns, results are presented by lipid class.

#### 1. Fatty Acids (FAs) and Fatty acid-derived oxylipins

Fatty acids undergo extensive remodeling across tissues, reflecting changes in lipid precursor availability (fig. 4 & 5). In the brain, LL treatment reduced DGLA, DPA(n-3/n-6) and DHA in the right hemisphere, with sex-specific responses: DHA decreased in males and DGLA in females. Here, the oxylipin 12-KETE was broadly downregulated across sexes, while 15-HETrE decreased only in the merged group. In the hypothalamus, AA was the only fatty acid found to be increased in females, where sex-dependent modulation of oxylipins was also observed: females upregulated AA-derived mediators (5-HETE, 5-KETE, 8-HETE, 15-HETE, 5,15-DiHETE and15-KETE) and DHA-derived oxylipins (4-HDHA, 17-HDHA, 14-HDHA + 7-HDHA), whereas males showed reductions in 12-KETE. Similarly, 12-HETE and 12-KETE lipids were decreased in the male midbrain along with 12-HEPE, whereas only females increased 4-HDHA.

**Figure 4.**
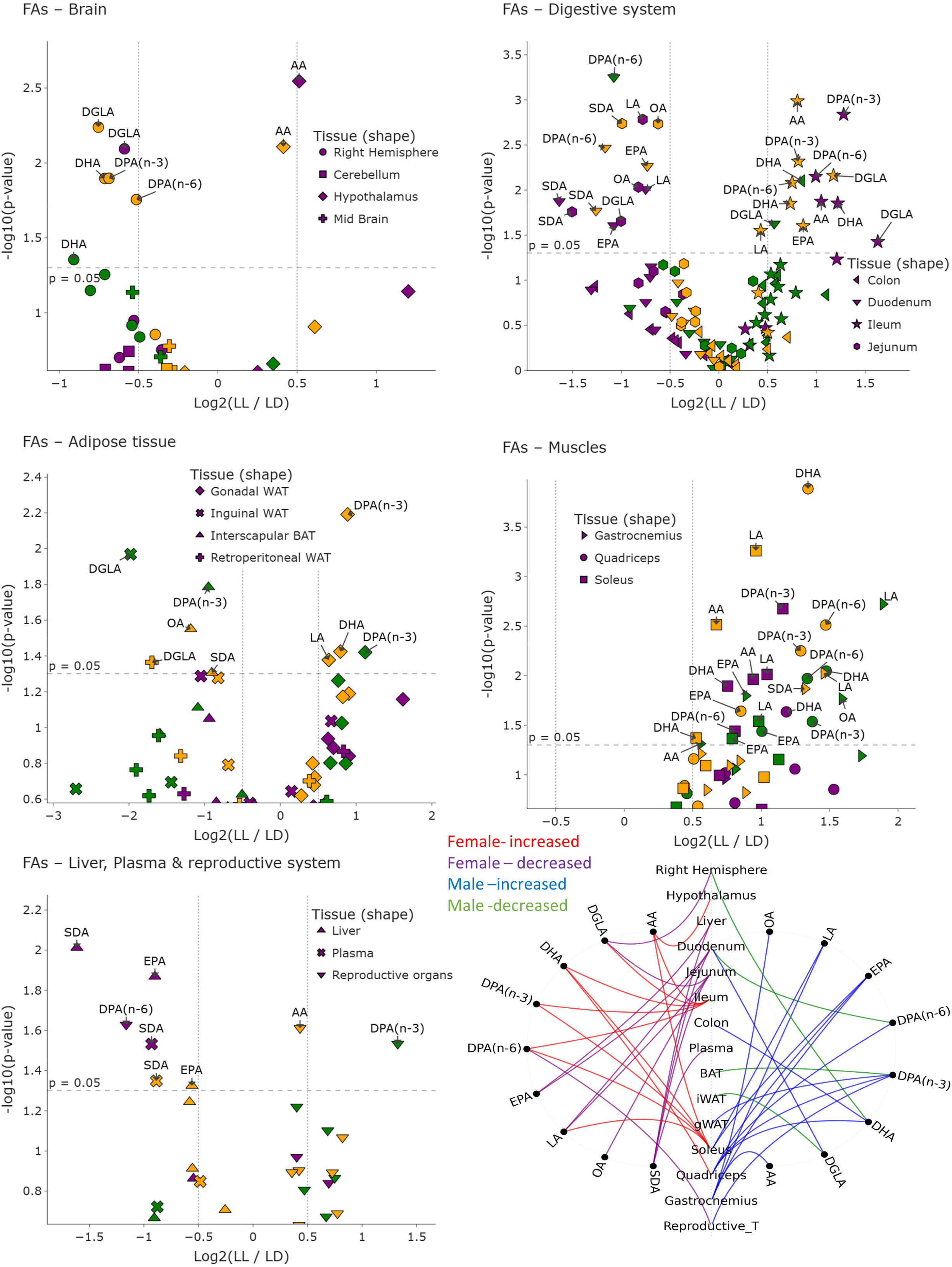
volcano plot of FAs; volcano plot showing differential levels of fatty acid (FAs) between LL and LD across all tissues analyzed. Each point represents one FAs measured in a specific tissue, with symbol shapes indicating the corresponding organ. Data are stratified by sex: females (purple), males (green), and combined (merged sex; yellow). The x-axis shows the log□ fold change (LL/LD) and the y-axis shows the –log□□(p-value). Molecules above the dashed horizontal line reached nominal statistical significance (p < 0.05). Labels highlight representative significant lipid mediators in each tissue. Chord Diagram (summary figure of FA changes; bottom right); Significant fatty acid changes across tissues in female and male mice under constant light exposure. Tissue nodes are aligned vertically, molecules appear on sex-specific arcs, and curves indicate LL-vs-LD regulation colored by sex and direction.

**Figure 5.**
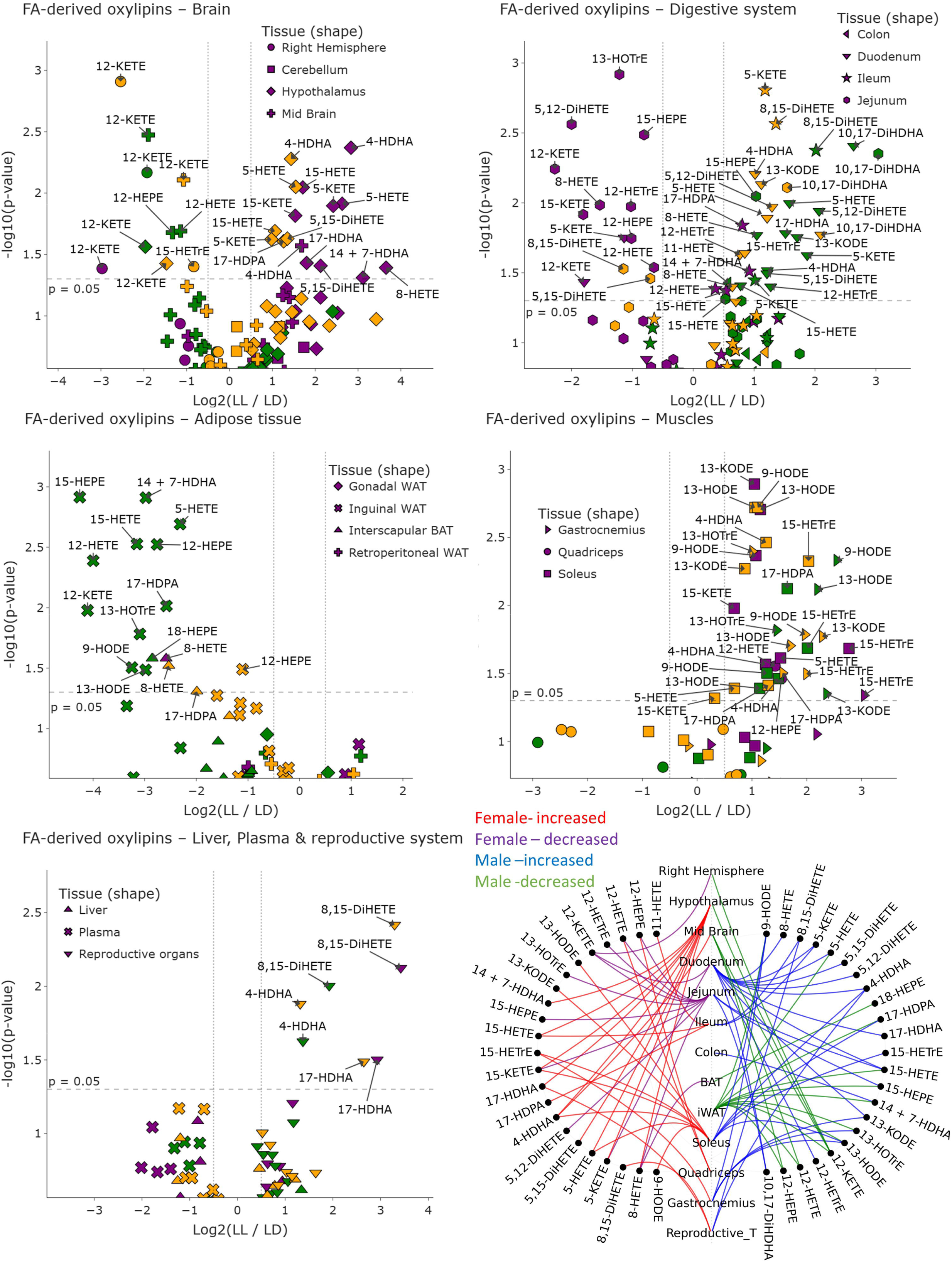
volcano plot of FA-derived Oxylipins; volcano plot showing differential levels of Fatty acid (FAs) derived oxylipins between LL and LD across all tissues analyzed. Each point represents one FAs-oxylipins measured in a specific tissue, with symbol shapes indicating the corresponding organ. Data are stratified by sex: females (purple), males (green), and combined (merged sex; yellow). The x-axis shows the log□ fold change (LL/LD) and the y-axis shows the –log□□(p-value). Molecules above the dashed horizontal line reached nominal statistical significance (p < 0.05). Labels highlight representative significant lipid mediators in each tissue. Chord Diagram (summary figure- of FA derived oxylipin changes; bottom right); Significant FA derived oxylipin changes across tissues in female and male mice under constant light exposure. Tissue nodes are aligned vertically, molecules appear on sex-specific arcs, and curves indicate LL-vs-LD regulation colored by sex and direction.

In the gastrointestinal tract, LL caused segment-specific effects that were almost exclusively observed in females or when the sexes were merged for fatty acids: in the duodenum, the n-3FAs SDA and DPA were reduced along with the n-6 FAs LA and DPA(n-6), with LA and SDA similarly reduced in the jejunum along with OA. In marked contrast the ileum saw increases in n-3 (EPA, DPA(n-3), DHA) and n-6 DGLA, AA, DPA(n-6)) FAs while no changes were observed in the colon other than an increase in DHA in males. Interestingly, several duodenal LA-, DGLA-, AA- and DHA-derived oxylipins (e.g 13-KODE, 15-HETrE, 5-HETE, 5-KETE, 12-HETrE, 4-HDHA, 10,17-DiHDHA) were upregulated in males and when sexes were combined. In marked contrast few changes in the jejunum were observed in males, while in females oxylipins (e.ge 13-HOTrE, 15-HETE, 12-HEPE) were decreased in this section of the small intestine. Strong changes in the Ileum were limited to increases in 5-KETE, 8,15-DiHETE and 10,17-DiDHA, largely driven by changes in males. and increases in DHA-derived 10,17-DiHDHA, along with sex-specific oxylipin changes. In the colon, similar to FAs, oxylipin levels were largely unchanged, except for in males that showed increased 14 + 7-HDHA, a DHA-derived oxylipin.

Shifting to peripheral metabolic tissues, Adipose depots exhibited heterogeneous responses to LL. In gonadal WAT, LL increased DPA(n-3), DHA, and LA, although the rise in DPA(n-3) appears to be driven by an increase in males. Conversely, in other adipose tissues, changes occurred in the opposite direction. In inguinal WAT and retroperitoneal WAT, LL decreased DGLA, an effect restricted to males, which also showed broad downregulation of oxylipins derived from LA (9-HODE, 13-HODE), AA (5-HETE, 15-HETE, 12-HETE, 12-KETE), ALA (13-HOTrE), EPA (15-HEPE, 12-HEPE), DPA (17-HDPA) and DHA- (14+7-HDHA). Among these only 12-HEPE was also decreased in the sex-merged group. Unlike WAT depots, BAT showed a more restricted pattern of downregulation consisting of SDA and OA, along with a decrease in DPA(n-3). Here, males exhibited decreased 18-HEPE, whereas females showed downregulation of 8-HETE, resulting in its reduction also in the merged sex group.

In contrast to adipose tissue, the muscle tissues (quadriceps, gastrocnemius, and soleus) showed robust increases in fatty acid and fatty acid derived oxylipins. In the gastrocnemius, when combining the sexes LL increased LA and SDA, along with elevations across several oxylipins, including LA-derived 9-HODE, 13-KODE, 13-HODE, ALA-derived 13-HOTrE, AA-derived mediator 15-HETrE and DPA-derived 17-HDPAe. In this muscle group the males generally drove these changes except for 15-HETrE which was increased in females. In the quadriceps, LL elevated DHA, DPA (n-3 and n-6), and EPA, again driven largely by increases in males, though DHA also increased in females. These changes were accompanied by elevated levels of 12-HETE and 12-HEPE in females. FA and oxylipin changes in the soleus were instead was more evident in females, with LL increasing LA, AA, and DHA and DPA (n-3 and n-6), though males showed increases in LA and EPA. Notably, FA derived oxylipins levels have increases similar to their precursors, for example in females we observed increased levels of LA-derived 13-KODE, 13-HODE, 9-HODE; AA-derived 15-KETE, 15-HETrE, 5-HETE; and the DHA-derived 4-HDHA. Similarly, in males, LL increased LA-derived 9-HODE and 13-HODE, AA-derived 15-HETrE, DPA-derived 17-HDPA, and DHA-derived 4-HDHA. All of these elevations were also reflected in the merged dataset.

Despite the widespread upregulation of fatty acids and oxylipins in skeletal muscle, the liver displayed a contrasting response, with reduced EPA in general and SDA only in females. SDA similarly declined in the plasma under LL, particularly in females. No changes in oxylipin levels were observed in either the liver or plasma. Outside somatic tissues, in the reproductive system, the female ovary showed a decrease in DPA(n-6) along with increases in AA-derived 8,15-DiHETE and DHA-derived 17-HDHA. In contrast, male reproductive tissue exhibited increased DPA(n-3) along with elevations in AA-derived 8,15-DiHETE and DHA-derived 4-HDHA. These oxylipin patterns persisted in the merged dataset, which showed consistent upregulation of 8,15-DiHETE, 4-HDHA, and 17-HDHA, highlighting a clear sex-dependent conservation in long-chain PUFA conversion to oxylipins.

#### 2. *N*-Acylethanolamines and NAE-derived oxylipins

NAEs showed tissue-specific regulation under LL exposure (Fig. 6 & 8). In the brain, regions like the cerebellum exhibited increased levels of PEA, OEA, AEA and DHEA, especially in females, with notable elevations in PEA, SEA, OEA, AEA and DHEA. Males, however, showed minimal response, limited to an increase in 17-HDPA-EA. The hypothalamus in females also showed significant increases in LEA and AEA, with a decrease in SEA, indicating sex-specific eCBome responses. Conversely, the midbrain displayed downregulation of PEA, SEA and EPEA, suggesting reduced NAE signaling. In the right hemisphere, LL exposure increased LEA-derived 13-HODE-EA in males, but not in females or the merged dataset. Outside the CNS, LL generally reduced NAEs in the gastrointestinal tract, with specific decreases in OEA, LEA and EPEA in the duodenum, and reductions in SDEA in the jejunum. The ileum showed decreased OEA and LEA, primarily in males, while the colon exhibited decreased LEA and 13-HODE-EA, especially in males. In adipose tissues, LL increased PEA and OEA in gonadal WAT, with females showing broader responses, including increases in PEA and DPEA, and a decrease in EPEA. In inguinal WAT, LL elevated EPEA in both sexes, but males also showed reductions in SEA and 13-HODE-EA. In interscapular BAT, LL decreased EPEA in females. Skeletal muscles were largely unresponsive, except for LEA increases in female soleus and quadriceps. The liver showed consistent downregulation of NAEs, with reductions in LEA and DHEA. In the reproductive system, the testis showed increased OEA in males, indicating sex specific NAE regulation.

**Figure 6.**
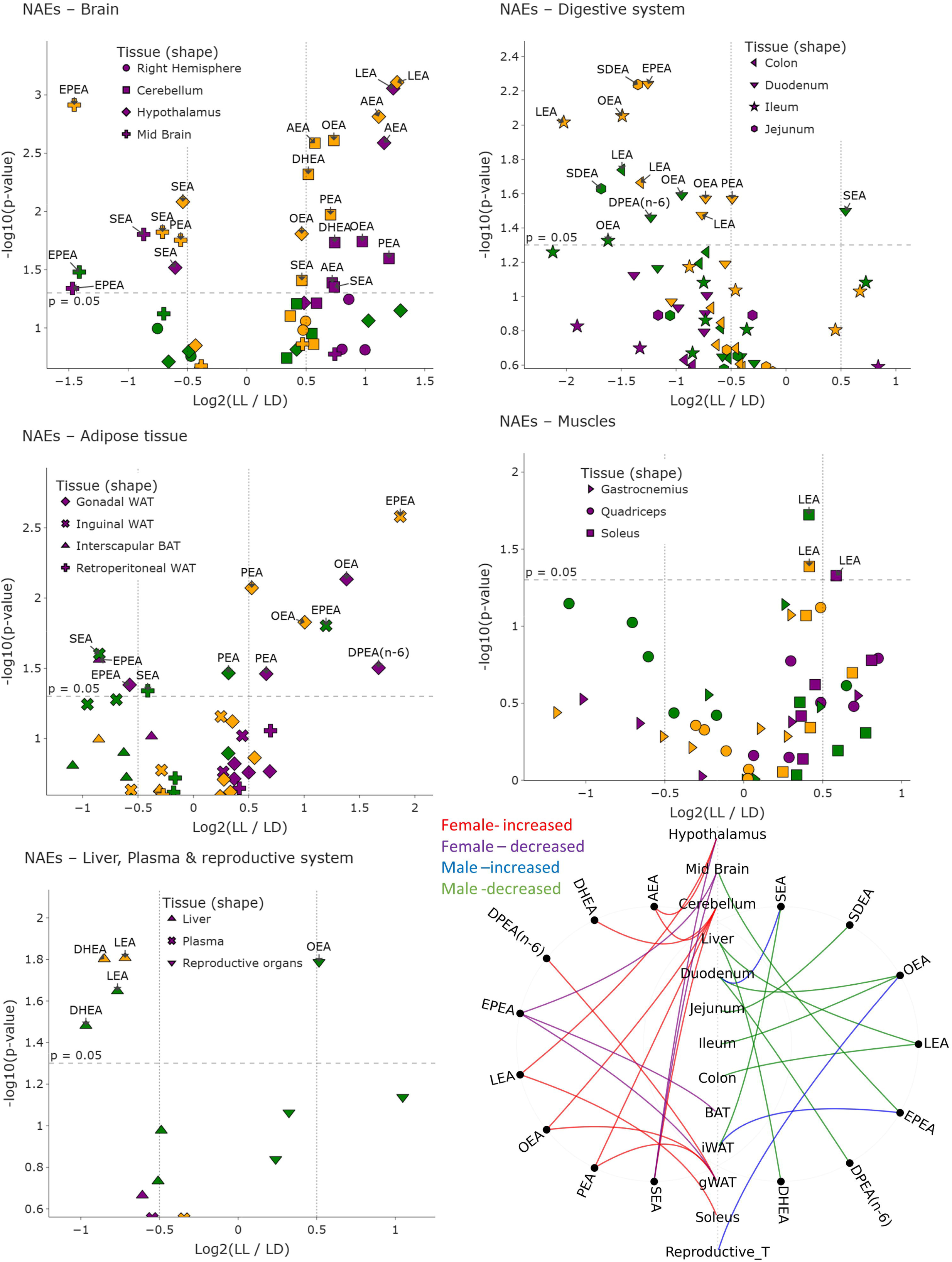
volcano plot of NAEs; volcano plot showing differential levels of N-acylethanolamines (NAEs) between LL and LD across all tissues analyzed. Each point represents one NAE measured in a specific tissue, with symbol shapes indicating the corresponding organ. Data are stratified by sex: females (purple), males (green), and combined (merged sexes; yellow). The x-axis shows the log□ fold change (LL/LD) and the y-axis shows the –log□□(p-value). Molecules above the dashed horizontal line reached nominal statistical significance (p < 0.05). Labels highlight representative significant lipid mediators in each tissue. Chord Diagram (summary figure of NAE changes; bottom right); Significant NAEs changes across tissues in female and male mice under constant light exposure. Tissue nodes are aligned vertically, molecules appear on sex-specific arcs, and curves indicate LL-vs-LD regulation colored by sex and direction.

**Figure 7.**
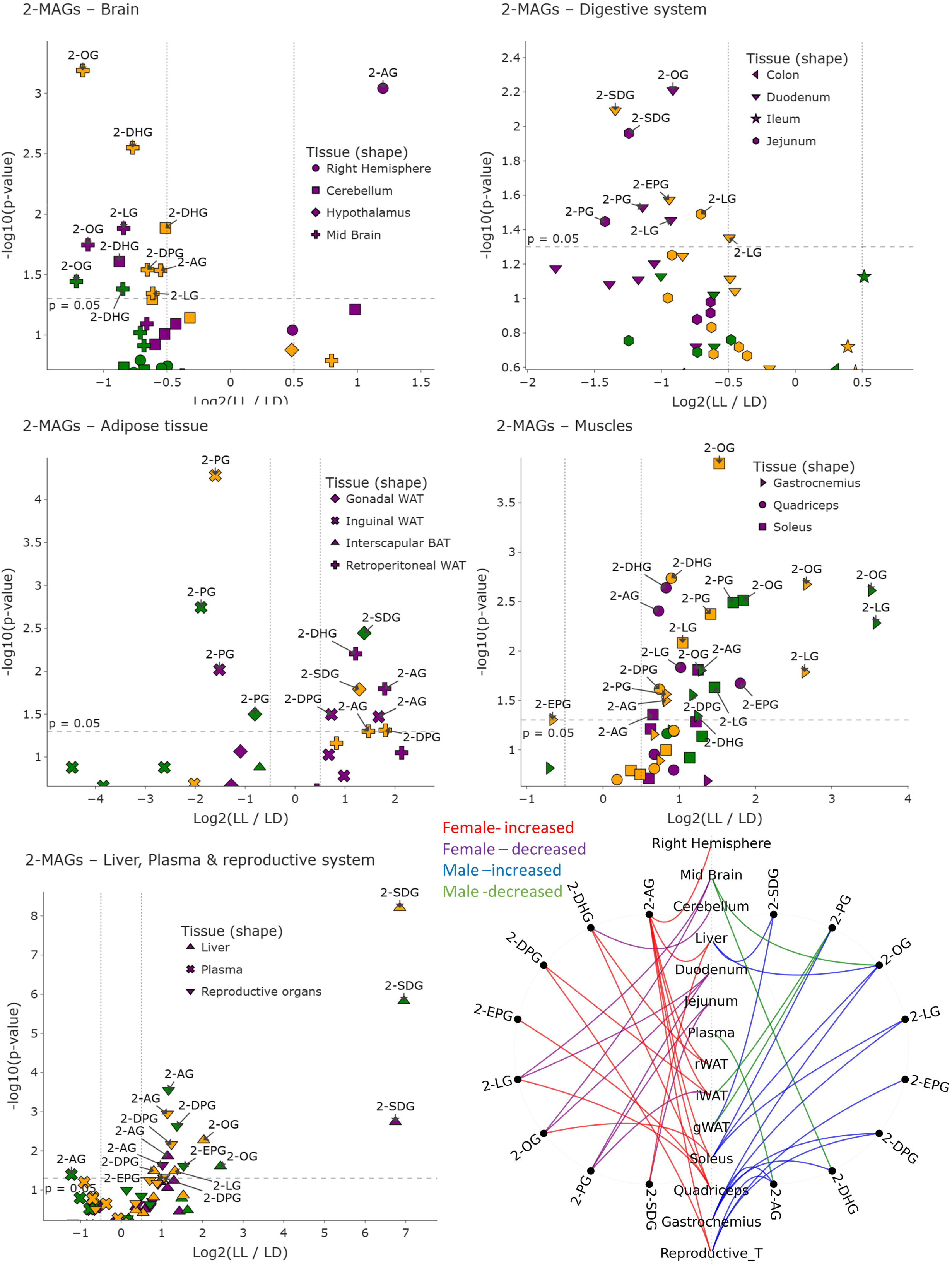
volcano plot of 2-MAGs; volcano plot of monoacylglycerols (2-MAGs) comparing LL and LD across all tissues. Each point corresponds to a MAGs species measured in one tissue, with symbol shapes indicating organ identity. Data are stratified by sex: females (purple), males (green), and combined (merged sexes; (yellow). The x-axis shows the log□ fold change (LL/LD) and the y-axis shows the – log□□(p-value). Molecules above the dashed horizontal line reached nominal statistical significance (p < 0.05). Labels highlight representative significant lipid mediators in each tissue. Chord Diagram (summary figure of 2-MAG changes; bottom right); Significant 2-MAGs changes across tissues in female and male mice under constant light exposure. Tissue nodes are aligned vertically, molecules appear on sex-specific arcs, and curves indicate LL-vs-LD regulation colored by sex and direction.

**Figure 8.**
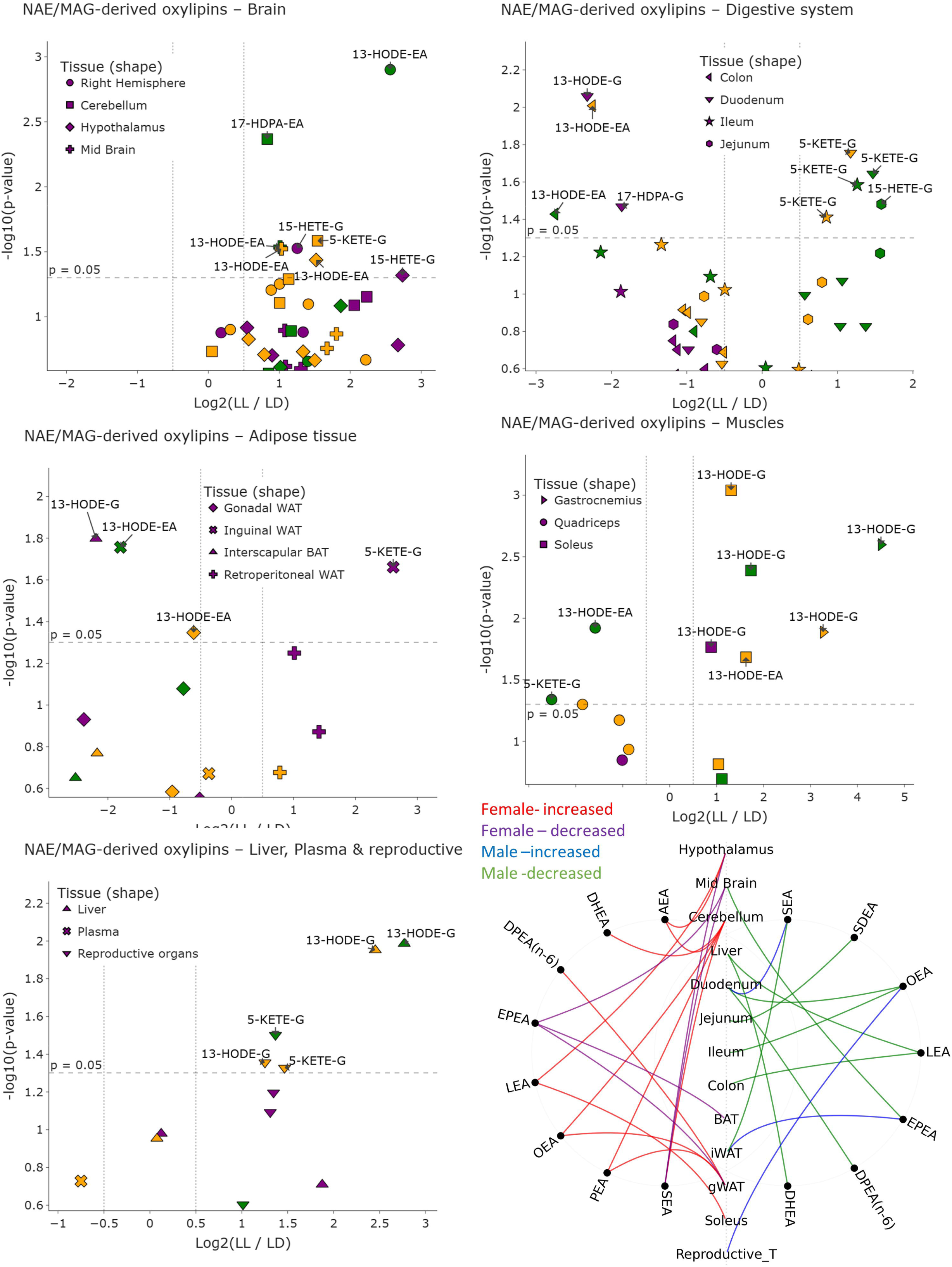
volcano plot of NAE- and 2-MAG-derived Oxylipins; volcano plot showing differential levels of N-Acylethanolamines (NAEs) and Monoacylglycerols (MAGs) derived Oxylipins between LL and LD across all tissues analyzed. Each point represents one NAES and MAGs-oxylipins measured in a specific tissue, with symbol shapes indicating the corresponding organ. Data are stratified by sex: females (purple), males (green), and combined (merged sexes; yellow). The x-axis shows the log□ fold change (LL/LD) and the y-axis shows the –log□□(p-value). Molecules above the dashed horizontal line reached nominal statistical significance (p < 0.05). Labels highlight representative significant lipid mediators in each tissue. Chord Diagram (summary figure of NAE- and 2-MAG-derived oxylipin changes; bottom right); Significant NAE- and 2-MAG-oxylipin changes across tissues in female and male mice under constant light exposure. Tissue nodes are aligned vertically, molecules appear on sex-specific arcs, and curves indicate LL-vs-LD regulation colored by sex and direction.

#### 3. Monoacylglycerols and 2-MAGs-derived oxylipins

Metabolic remodeling in neural, metabolic, and peripheral tissues was observed following specific characteristics (fig 7 & 8). In the midbrain, reductions in several 2-MAGs such as 2-OG, 2-LG, 2-AG, 2-DPG and 2-DHG were noted, with sex-specific patterns: males showed decreases in 2-OG and 2-DHG, while females exhibited reductions in 2-LG and 2-OG. These changes suggest increased turnover of 2-MAGs in this brain region. Additionally, in the female cerebellum, 2-DHG decreased, whereas 2-AG increased in the right hemisphere, coinciding with elevated levels of its oxylipin product, 15-HETE-G, which was upregulated in both the female right hemisphere and hypothalamus. In the gastrointestinal tract, particularly the small intestine, reductions in 2-MAGs such as 2-SDG and 2-EPG were observed, with females showing broader suppression including 2-PG, 2-OG, and 2-LG. These decreases were accompanied by reductions in oxylipins like 13-HODE-G and 17-HDPA-G. Conversely, in males, 2-AG derived HETE- and KETE-glycerols increased. Along the intestinal tract, the jejunum showed decreased 2-LG, with females also exhibiting reductions in 2-SDG and 2-PG. Adipose tissues displayed mixed responses, with some MAGs increasing and others decreasing. In gonadal white adipose tissue (WAT), 2-PG decreased in males, while 2-SDG increased in both sexes. In inguinal WAT, 2-PG decreased across groups, but females showed increases in 2-DPG and 2-AG, along with elevated 2-AG derived oxylipin 5-KETE-G. Retroperitoneal WAT exhibited increased 2-DHG and 2-AG in females, indicating broader MAG elevation. Skeletal muscles, including gastrocnemius, quadriceps, and soleus, showed significant sex-specific increases in MAGs, with notable elevations in 2-OG, 2-LG, 2-AG, and 2-DPG, and associated oxylipins like 13-HODE-G. Hepatic tissue demonstrated a consistent increase in multiple MAGs, with females showing elevated 2-SDG and 2-AG, and males showing increases in 2-SDG and 2-OG. The reproductive organs also showed upregulation of 2-AG and 2-DPG, with males additionally increasing 2-EPG. Systemically, plasma MAGs showed a decrease in 2-AG only in males, indicating tissue-specific and sex-specific metabolic adaptations.

#### 4. Prostaglandins (PGs) and Eicosanoid-Derived Lipid Mediators

Prostaglandins exhibit significant tissue-specific and sex-dependent remodeling (fig 9). In the brain, the cerebellum shows increased PGB[, especially in females, along with PGE[and 1a,1b-dihomo-PGF[α. The hypothalamus displays elevated PGD[in both sexes, sexes, with TXB[specifically increased in males; coordinated increases in PGD[, TXB[, PGE[, and PGE[are observed in the merged dataset. The midbrain shows a reduction in PGD[-EA, and the right hemisphere exhibits decreased 12-oxo-LTB[in males and overall. The duodenum demonstrates male-biased prostaglandin upregulation, including LTB[, 12-oxo-LTB[, PGB[, and 5,6-DiHETE, with coordinated increases in these mediators and a decrease in 12-HHTrE. The ileum shows increased PGF[_α_ in females, while PGF_3α_ is up in the merged dataset. In adipose tissue, PG levels vary: 6-keto-PGF[_α_ decreases in males, and PGE[-G increases in females and males. The jejunum displays decreased 17-oxo-DHA in females and increased PGE[-G, with males showing elevated PGF[α-EA. The colon shows increased 17-oxo-DHA and PGF[_α_ in males. Skeletal muscle exhibits sex-specific changes: in gastrocnemius, PGE[-EA decreases in females; in quadriceps, LL increases 12-HHTrE in females, while PGE[decreases in males; and in soleus, PGE[, PGD[, and PGF[_α_ are upregulated, especially in females. Liver responses are characterized by suppression of prostanoids like PGF[_α_ and PGE[, as well 12-HHTrE, mainly in males. Adipose depots show increased PGD[and PGB[in retroperitoneal and gonadal WAT, while BAT shows decreased PGF[_α_ and PGE[. Systemically, plasma 12-HHTrE and PGD[are reduced, with increased 12-oxo-LTB[in females and elevated 17-oxo-DHA, 12-oxo-LTB[, and PGE[in males.

**Figure 9.**
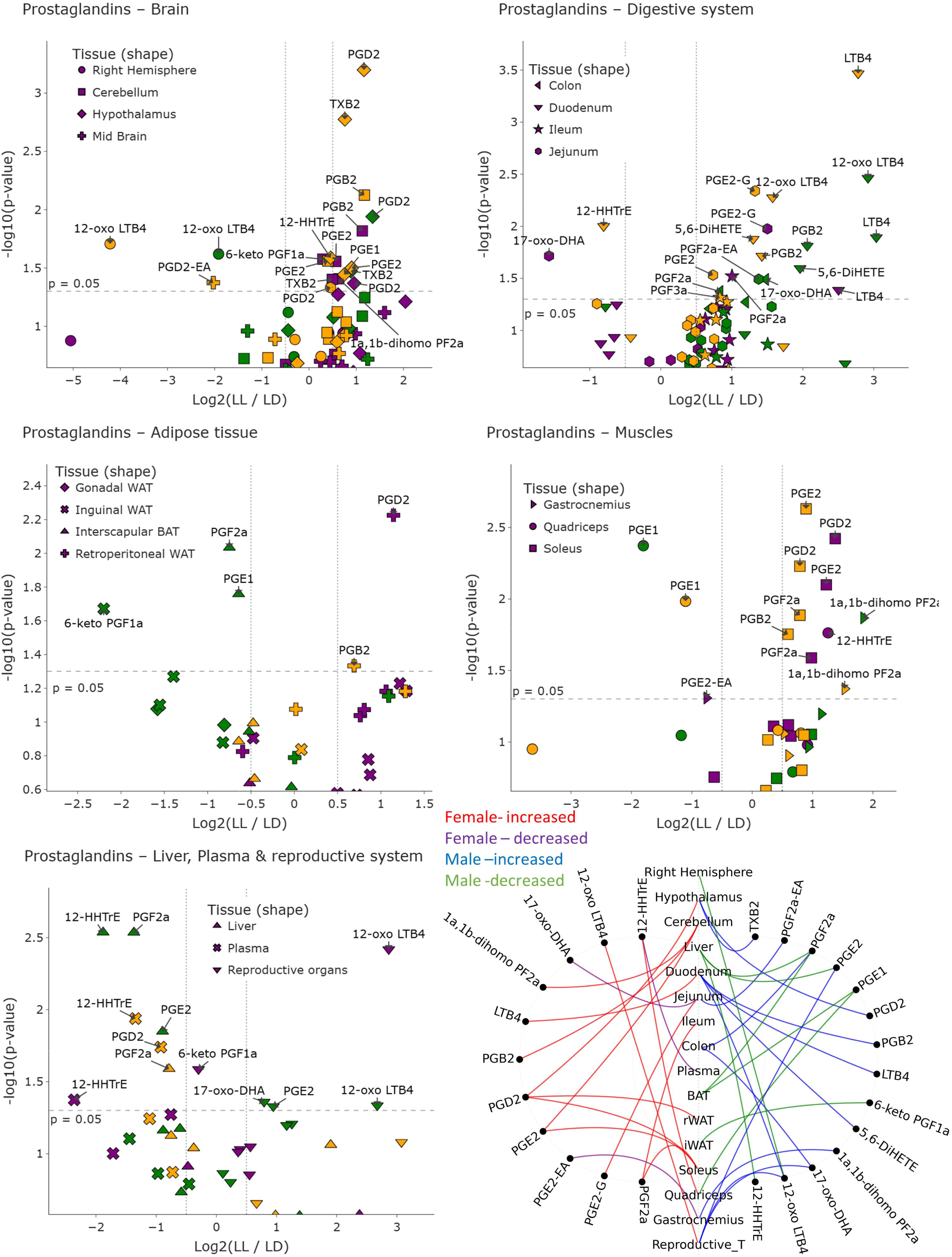
Volcano plot of PGs; volcano plot showing differential levels of prostaglandin (PG) between LL and LD across all tissues analyzed. Each point represents one PG measured in a specific tissue, with symbol shapes indicating the corresponding organ. Data are stratified by sex: females (purple), males (green), and combined (merged sexes; yellow). The x-axis shows the log□ fold change (LL/LD) and the y-axis shows the –log□□(p-value). Molecules above the dashed horizontal line reached nominal statistical significance (p < 0.05). Labels highlight representative significant lipid mediators in each tissue. Chord Diagram (summary figure of PG changes; bottom right); Significant PGs changes across tissues in female and male mice under constant light exposure. Tissue nodes are aligned vertically, molecules appear on sex-specific arcs, and curves indicate LL-vs-LD regulation colored by sex and direction.

Cytokine analysis reveals sex-specific immune responses (fig 10): females show increased IL-5, MIP-2, and IFN-γ, while males exhibit elevated IL-2 and trends toward increased IFN-γ. When combined, IFN-γ, IL-1β, IL-13, and GM-CSF are elevated, indicating systemic inflammatory modulation aligned with female-driven lipid remodeling.

**Figure 10.**
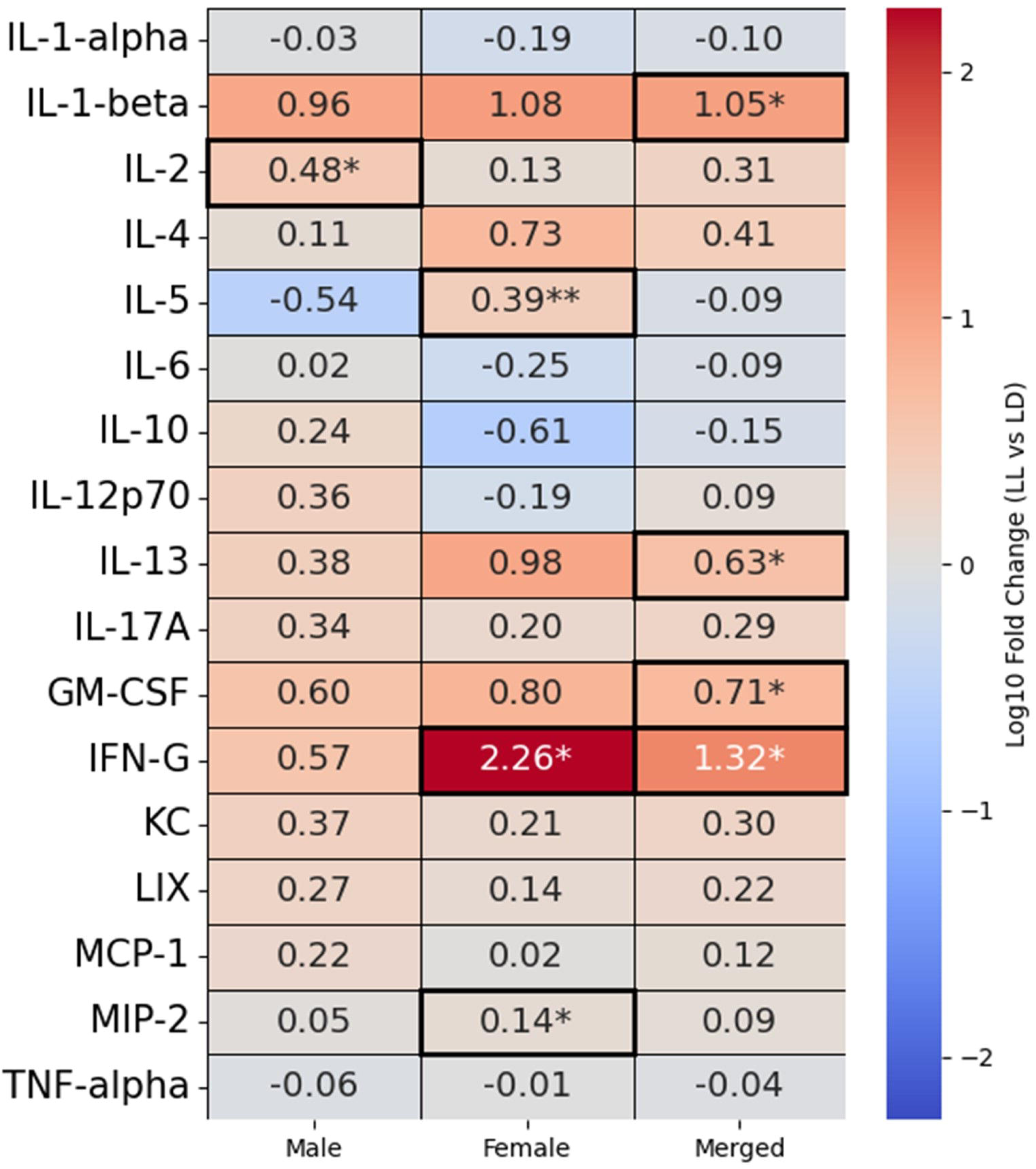
Heatmap of plasma cytokine responses under LL) LD exposure. Shown are log□□ fold changes (LL vs LD) for individual cytokines in males, females, and merged data. Color coding indicates the magnitude and direction of fold change (red = increase, blue = decrease). Notably, IL-1β, IL-2, IL-13, GM-CSF, and IFN-γ were significantly increased, with sex-specific patterns including elevated IFN-γ, MIP-2 and IL-5 in females and increased IL-2 in males. Significant changes are highlighted with asterisks (* for p < 0.05, ** for p < 0.01).

## Discussion

Circadian rhythm disruption profoundly alters host physiology and increases susceptibility to metabolic and inflammatory conditions (4). Given the intimate role of gut microbiota and the eCBome in regulating multiple aspects of metabolic health (47, 61) and responding to circadian cycling (26, 50), we went on to determine how circadian rhythm disruption using constant light modulates eCBome lipid levels and gut bacteria. We selected ZT11 and CT11 for organ collection because it represents a transition point into the dark cycle and active phase, making it ideal for studying metabolic and circadian-regulated processes (27). Our findings reveal clear sex-dependent adaptive strategies to constant light (LL) with respect to changes induced in bioactive lipid levels in different organs and fecal bacteria composition.

LL exposure altered the gut microbiota in a sex-dependent manner. In females, taxa such as *Rikenellaceae, Butyricicoccaceae*, *Alistipes*, *Eisenbergiella*, and *Enterorhabdus* were enriched, while *Erysipelotrichaceae*, *Turicibacter*, and *Ruminococcaceae UCG-005* were depleted. Males showed fewer compositional shifts, with increases in *Tuzzerella* and *Oscillospirales*-*UCG-010* and reductions in *Ruminococcaceae*. Several of these taxa have been linked to circadian disruption or rhythmicity in prior studies. For example, in constant darkness (DD), *Alistipes* (62) significantly changes in the opposite direction to our protocol (constant light), whereas *Rikenellaceae* (63) increased in the same direction. These differences likely reflect the distinct circadian stressors of LL versus DD (loss of light–dark cycles vs. loss of photic cues), highlighting their sensitivity to circadian misalignment.

Indeed intestinal epithelium disruption of *Bmal1* leading to disruption of the intestinal clock disrupts *Alistipes* rhythmicity (64), and constant light exposure causes the opposite effect on its levels compared to what we report here (64).Thus our data suggests that this genus may be particularly sensitive to light exposure, though the potential effects of this would be dependent on the species affected, as different *Alistipes spp.* have been found to associate with positive or negative physiological effects (65). The consistent reduction of *Turicibacter*, a genus positively associated with metabolic health, potentially by modifying bile acids and thus affecting lipid metabolism (66), further suggests that LL disrupts microbial taxa normally linked to rhythmicity and beneficial functions.

Despite pronounced taxonomic alterations in females, SCFA production increased only in males, with higher levels of isobutyric, butyric, isovaleric, and valeric acids. These shifts coincided with reductions in the *Ruminococcaceae* family and enrichment of *Oscillospirales-UCG-010* and *Tuzzerella* at the genus level. The *Oscillospirales*-*UCG-010* correlate with SCFA levels (67). This enrichment is consistent with the elevated fecal SCFA levels observed in males, suggesting a functional link between microbial composition and metabolite output. *Tuzzerella*, although not a major SCFA producer, has been reported to increase in models of gut inflammation (68), where it modulates immune responses by regulating T cell infiltration (69). Its enrichment under LL may therefore reflect proliferation during inflammatory stress. In parallel, SCFAs such as acetate, propionate, and butyrate can enter the liver and directly modulate circadian clock gene expression (51). SCFAs also exert local benefits, including reducing systemic inflammation, improving hepatic lipid metabolism, and ameliorating mucosal injury in the small intestine (70). Given established associations between SCFAs and sleep duration (71), their elevation in males may indicate a behavioral or metabolic protective adaptation to LL-induced circadian rhythm disruption. These elevated levels of SCFAs might potentially have affected the levels of oxylipins within the distal intestinal tract, which in fact increased in the duodenum and jejunum of male LL mice, and decreased in females. Indeed, in comparison to the duodenum, the lower intestinal regions showed also significant increases in long-chain PUFAs, including DHA and its LOX-derived oxylipins, accompanied by suppression of NAEs and prostaglandins, emphasizing the distal gut as a hotspot of circadian stress responses.

In females, although the microbial community composition changed markedly, SCFA levels did not. Instead, intestinal lipid remodeling reflected widespread depletion of protective mediators. The proximal small intestine showed downregulation of fatty acids (LA, OA), MAGs (2-SDG), and prostaglandins (PGE_2_). NAEs were largely unchanged in the duodenum and jejunum but markedly suppressed in the ileum (LEA, PEA). While DHA was increased in the ileum where, unlike males, its oxidized metabolites were not. These signatures align with prior reports linking circadian disruption to intestinal inflammation (72) and heightened metabolic disease susceptibility (73, 74). Such sex-specific differences may reflect greater disruption of intestinal clock gene expression and metabolic adaptation in females (75) and their propensity of female mice to lose weight under protocols of light-cycle disruption while males gain weight (76).

Under LL, females showed suppression of lipid mediators in the proximal intestine but activation in the distal gut, muscle, and reproductive organs (e.g., DPA n-3, 2-AG, 15-HpEDE, PGD[), indicating a redistribution of lipid signaling from the gut toward peripheral tissues. Females engaged NAE-centered eCBome signaling (e.g., AEA, OEA, PEA) in the brain and liver, accompanied by systemic inflammatory cytokine responses. In contrast, males exhibited activation of oxylipin and 2-MAG pathways in the intestine and liver (e.g., 5-HETE, 5-KETE, 10,17-DiHDHA, 14-/ 7-HDHA, 2-AG) and in skeletal muscle (e.g., 2-AG, 2-LG). In the adipose tissues, particularly inguinal WAT, males showed coordinated suppression of anti-inflammatory oxylipins (e.g., 15-HEPE, 5-/ 15-HETE, 14-/ 7-HDHA, 13-HODE-G, 6-keto PGF[α). Overall, males relied on SCFA- and ω-3-derived resolution pathways, particularly in the colon and liver, whereas females engaged eCBome-driven signaling and immune activation under circadian disruption. The liver exhibited a mixed regulatory profile. In females, ω-3 fatty acids (SDA, EPA) decreased alongside increased 2-MAGs (2-SDG, 2-AG). In males, prostaglandins (PGF[α, PGE[) and NAEs (LEA, DHEA) were suppressed, while also in this sex 2-MAGs (2-SDG, 2-OG) increased. When sexes were combined, reduced prostaglandins and NAEs persisted and 2-MAGs remained elevated. This shift mirrors metabolic liver disease, where reduced ω-3 fatty acids (77) and prostaglandins (78) promote accumulation of 2-MAG-derived diacylglycerol (DAG) via monoacylglycerol acyltransferases (MGATs), subsequently forming triacylglycerols (TAGs) through DGAT2 (79), an enzyme that itself exhibits rhythmic expression (80). This pathway contributes to non-alcoholic fatty liver disease (NAFLD) (81). Interestingly, however, 2-SDG emerged as the most responsive hepatic molecule under LL in both sexes. This 2-MAG is derived from the ω-3 fatty acid, SDA, which is associated with potential anti-inflammatory properties (82). However, the precise biological role of 2-SDG remains to be clarified both metabolically and with respect to its potential involvement in light-cycle disruption effects on liver metabolism.

The brain showed a sex dependent response; in females, the cerebellum displayed robust upregulation of NAEs and prostaglandins which is consistent with neuroprotective response of NAEs (83–86). This response was absent in males. The hypothalamus also showed LOX-derived oxylipins (derived from both FAs and 2-MAGs; e.g., 12-HETE, 15-HETE, 15-HETE-G) proposed to modulate neuroinflammation and neuronal lipid signaling (87), reflecting the sensitivity of this brain area to circadian stress. Pharmacological CB_1_ activation reduces the phase-shifting effects of light on the suprachiasmatic nucleus in male but not female mice (88), further supporting tissue- and sex-specific differences in circadian responsiveness. By contrast, the midbrain and right hemisphere exhibited marked downregulation of NAEs, 2-MAGs, oxylipins, and long-chain fatty acids (e.g., DGLA, DHA, DPA), with males showing a more prostaglandin-centered response. Although NAEs are generally considered neuroprotective, multiple studies indicate they can also contribute to pathogenesis and elevate disease risk. Elevated NAE levels have been linked with aging, neuroinflammation, and ischemia (89), can directly trigger cell death (90, 91), and are increased in psychotic patients (92). This duality likely reflects the several molecular targets for each of these mediators, ranging from CB_1_ and TRPV1, which may play pro-inflammatory and brain function disrupting effects, to GPR110, PPARs and CB_2_, which instead play pro-homeostatic anti-inflammatory actions (37). Circadian disturbances are highly prevalent in schizophrenia and psychosis (93) and with aging, the amplitude of circadian rhythms progressively declines (94). Collectively, these findings suggest that female brains preferentially engage NAE pathways as a protective buffer against circadian rhythm disruption. However, given the dual role of NAEs in both protection and pathology, their alteration under light-cycle disruption may lead to maladaptive neurobiological changes in females, potentially contributing to greater vulnerability to shift-work disorder, stress, and insomnia (95).

LL exposure induced FA accumulation across several skeletal muscles, with little or no effect, however, on eCBome mediators. Sleep restriction and loss of circadian rhythmicity are known to cause atrophy and impaired metabolic plasticity in muscles such as the quadriceps, soleus, and gastrocnemius (96, 97). These muscles are enriched for rhythmic genes involved in fatty acid metabolism, mitochondrial oxidative phosphorylation, and glucose handling (98, 99). Moreover, post-flight biopsies of human soleus have demonstrated susceptibility to lipid accumulation under circadian misalignment (100). Within this context, our data show that LA and OA accumulated in the gastrocnemius, and LA in the soleus, indicating maladaptive lipid storage particularly within glycolytic muscle fibers, consistent with reduced oxidative flexibility and altered energy metabolism.

In contrast to muscles the adipose tissue was less effected by circadian rhythm disruption; in inguinal WAT, females showed increased 2-MAGs, including 2-AG, indicating enhanced cannabinoid receptor CB[activity (101). Adipose CB[signaling is known to impair insulin sensitivity, enhance lipogenesis, and suppress mitochondrial oxidative capacity, ultimately promoting lipid accumulation and endoplasmic reticulum stress (102, 103). These changes may therefore represent an early maladaptive metabolic response to circadian disruption. In contrast, males exhibited coordinated suppression of multiple oxylipins and NAEs, suggesting loss of local anti-inflammatory tone that may predispose to low-grade inflammation (104). In retroperitoneal WAT, increases in prostaglandins (PGD[, PGB[) and 2-MAGs (2-AG, 2-DPG, both of which have been shown to have context-dependent anti- and pro-inflammatory activities (43)) highlighting coordinated eCBome response within a given family of lipids. The decreased levels of DGLA may indicate limited substrate availability for the production of anti-inflammatory prostaglandin synthesis and thus a relative increase in pro-inflammatory ones, potentially exacerbating inflammation (105). Interscapular BAT displayed suppression of bioactive lipids including ω-3 fatty acids, EPEA, and prostaglandins, consistent with reduced lipid signaling capacity that could blunt thermogenic and oxidative functions (106). An increase in 2-MAGs across multiple adipose tissues may contribute to metabolic dysfunction and obesity-related complications, such as insulin resistance and type 2 diabetes, through their role as intermediates in triglyceride synthesis (107). Together, these findings align with the LL disruption of adipose lipolysis and its coordination with skeletal muscle fatty acid uptake to cause a systemic metabolic dysfunction (108).

Reproductive tissues displayed a similar remodeling under constant light with having increased the 2-MAGs, and FA-oxylipins in male and female. In particular, the males showed upregulation of OEA, DPA(n-3), PGE[, multiple oxylipins (4-HDHA, 17-oxo-DHA, 5-KETE-G), and 2-MAGs (2-AG, 2-DPG, 2-EPG). In females, we also observed increased in oxylipins (12-oxo-LTB4, 8,15-DiHETE, 17-HDHA), and 2-MAGs (2-AG, 2-DPG) were elevated while DPA(n-6) was downregulated. Both testes and ovaries have functional molecular clocks; in males, these clocks regulate testosterone production, spermatogenesis, and sperm motility (109), and in females, they coordinate hypothalamic-pituitary-gonadal (HPG) axis output with ovarian steroidogenesis and ovulation timing, partly through rhythmic prostaglandin and NAE signaling (110). Disruption of these rhythms is therefore likely to impair fertility. Clinical observations support this interpretation. Elevated seminal 2-AG has been associated with reduced sperm motility in men (111), while high 2-AG levels are reported in women with endometriosis and infertility (112). Similarly, increased oxylipins and associated oxidative stress can impair gamete function and reproductive outcomes (113), linking circadian disruption, dysregulated lipid signaling, and chronic inflammation to infertility. Supporting this link, clinical studies indicate that misalignment of circadian rhythms such as that caused by shift work or constant light, leads to reduced fertility, irregular cycles, and lower conception rates (114).

Systemic cytokine profiling revealed distinct sex-specific immune signatures under constant light. Females displayed elevated IL-5, IFN-γ, and the neutrophil-recruiting chemokine MIP-2, whereas males showed only IL-2 significant increase. This pattern suggests that circadian disruption amplifies immune responsiveness more strongly in females and is consistent with computational and experimental models indicating that circadian clocks modulate immune activity in a sex-dependent manner (115). IFN-γ levels were reported to be significantly altered in Per2 mutant mice, which lacked a daily rhythm (116), and studies in non-human primates showed that the severity of circadian rhythm disturbances correlates with plasma IFN-γ levels (117). Interestingly, despite the heightened cytokine response, plasma lipid mediators showed a predominantly suppressive profile under constant light. In females, both SDA and the oxylipin 12-HHTrE were reduced, while males exhibited lower 2-AG levels.

The present study has some limitations. The relatively small sample size (n=4 per sex for each treatment) may have limited statistical power. We assessed tissues at a single circadian timepoint, which may not capture dynamic oscillations of microbial and lipid networks. Finally, the sex hormone status in females was not considered and could have contributed to variability in circadian responses.

### Conclusions

Constant light exposure disrupted circadian homeostasis, inducing sex-dependent alterations in fecal microbiota, eCBome and other bioactive lipids, and systemic immune networks. Females exhibited marked microbiota remodeling, depletion of protective intestinal lipids, and of NAE-mediated response in the brain area, accompanied by elevated pro-inflammatory cytokines such as IFN-γ, IL-5, and MIP-2. In contrast, males displayed increased fecal SCFAs and enhanced oxylipin levels in the intestine and liver, reflecting a metabolic adaptation centered on lipid oxidation rather than inflammation. The liver and reproductive tissues emerged as key sites of lipid mediator remodeling, with LL exposure often promoting ω-3 fatty acid depletion, 2-MAG accumulation, and prostaglandin and oxylipin pathway activation; patterns that are linked to metabolic stress and reproductive dysfunction. Collectively, these findings reveal that circadian disruption under constant light disturbs the coordination between microbial, lipidomics, and immune pathways in a tissue- and sex-specific manner, predisposing the organism to inflammation and metabolic imbalance.

## Abbreviations

LL: constant light
LD: light/dark
SCN: suprachiasmatic nucleus
SCFA: short-chain fatty acid
ECS: endocannabinoid system
eCBome: endocannabinoidome
NAE: *N*-acylethanolamine
2-MAG: 2-monoacylglycerol
2-AG: 2-arachidonoylglycerol
AEA: anandamide
PEA: *N*-palmitoyl-ethanolamine
OEA: *N*-oleoyl-ethanolamine
DHEA: N-docosahexaenoyl-ethanolamine
WAT: white adipose tissue
BAT: brown adipose tissue
DHA: docosahexaenoic acid
EPA: eicosapentaenoic acid
SDA: stearidonic acid
MASLD: metabolic dysfunction-associated steatotic liver disease
ZT / CT: zeitgeber time / circadian time
GC-FID: gas chromatography with flame ionization detection
LC-MS/MS: liquid chromatography–tandem mass spectrometry.

## Author contribution

PAP, performed experiments, analysis, and manuscript writing; TN, performed experiments, edited manuscript; TK and ED, analysis; NF, analysis and manuscript editing; VDM, analysis and manuscript editing; EM, experimental design and manuscript editing, and CS: experiment conceptualization, experimental design, manuscript editing and funding.

## References

1. Fagiani F, Di Marino D, Romagnoli A, Travelli C, Voltan D, Di Cesare Mannelli L, et al. Molecular regulations of circadian rhythm and implications for physiology and diseases. Signal Transduct Target Ther. 2022;7(1):41.

2. Michel S, Itri J, Han JH, Gniotczynski K, Colwell CS. Regulation of glutamatergic signalling by PACAP in the mammalian suprachiasmatic nucleus. BMC Neurosci. 2006;7:15.

3. Takahashi JS. Transcriptional architecture of the mammalian circadian clock. Nat Rev Genet. 2017;18(3):164–79.

4. Coomans CP, van den Berg SA, Houben T, van Klinken JB, van den Berg R, Pronk AC, et al. Detrimental effects of constant light exposure and high-fat diet on circadian energy metabolism and insulin sensitivity. FASEB J. 2013;27(4):1721–32.

5. Fonken LK, Workman JL, Walton JC, Weil ZM, Morris JS, Haim A, et al. Light at night increases body mass by shifting the time of food intake. Proc Natl Acad Sci U S A. 2010;107(43):18664–9.

6. Skinner NJ, Rizwan MZ, Grattan DR, Tups A. Chronic Light Cycle Disruption Alters Central Insulin and Leptin Signaling as well as Metabolic Markers in Male Mice. Endocrinology. 2019;160(10):2257–70.

7. Nelson RJ, Chbeir S. Dark matters: effects of light at night on metabolism. Proc Nutr Soc. 2018;77(3):223–9.

8. Parkes KR. Demographic and lifestyle predictors of body mass index among offshore oil industry workers: cross-sectional and longitudinal findings. Occup Med (Lond). 2003;53(3):213–21.

9. Boege HL, Bhatti MZ, St-Onge MP. Circadian rhythms and meal timing: impact on energy balance and body weight. Curr Opin Biotechnol. 2021;70:1–6.

10. Fonken LK, Nelson RJ. The effects of light at night on circadian clocks and metabolism. Endocr Rev. 2014;35(4):648–70.

11. Lemmer B, Oster H. The Role of Circadian Rhythms in the Hypertension of Diabetes Mellitus and the Metabolic Syndrome. Curr Hypertens Rep. 2018;20(5):43.

12. Gnocchi D, Custodero C, Sabba C, Mazzocca A. Circadian rhythms: a possible new player in non-alcoholic fatty liver disease pathophysiology. J Mol Med (Berl). 2019;97(6):741–59.

13. Zinna L, Verde L, Tolla MFD, Barrea L, Parascandolo A, D’Alterio F, et al. Chronodisruption enhances inflammatory cytokine release from visceral adipose tissue in obesity. J Transl Med. 2025;23(1):231.

14. Woodie LN, Johnson RM, Ahmed B, Fowler S, Haynes W, Carmona B, et al. Western diet-induced obesity disrupts the diurnal rhythmicity of hippocampal core clock gene expression in a mouse model. Brain Behav Immun. 2020;88:815–25.

15. Wang S, Bao Y, Wang L, Xie X, Lu Y. Association of dietary quality and dietary inflammatory potential with inflammatory markers: evidence from the national health and nutrition examination survey 2009-2018. Front Immunol. 2025;16:1596806.

16. Cermakian N, Westfall S, Kiessling S. Circadian clocks and inflammation: reciprocal regulation and shared mediators. Arch Immunol Ther Exp (Warsz). 2014;62(4):303–18.

17. Tsurudome Y, Yoshida Y, Hamamura K, Ogino T, Yasukochi S, Yasuo S, et al. Prostaglandin F2alpha Affects the Cycle of Clock Gene Expression and Mouse Behavior. International journal of molecular sciences. 2024;25(3).

18. Hablitz LM, Gunesch AN, Cravetchi O, Moldavan M, Allen CN. Cannabinoid Signaling Recruits Astrocytes to Modulate Presynaptic Function in the Suprachiasmatic Nucleus. eNeuro. 2020;7(1).

19. Sladek M, Liska K, Houdek P, Sumova A. Modulation of single cell circadian response to NMDA by diacylglycerol lipase inhibition reveals a role of endocannabinoids in light entrainment of the suprachiasmatic nucleus. Neuropharmacology. 2021;185:108455.

20. Hanlon EC. Impact of circadian rhythmicity and sleep restriction on circulating endocannabinoid (eCB) N-arachidonoylethanolamine (anandamide). Psychoneuroendocrinology. 2020;111:104471.

21. Acuna-Goycolea C, Obrietan K, van den Pol AN. Cannabinoids excite circadian clock neurons. J Neurosci. 2010;30(30):10061–6.

22. Sladek M, Sumova A. Modulation of NMDA-Mediated Clock Resetting in the Suprachiasmatic Nuclei of mPer2 (Luc) Mouse by Endocannabinoids. Front Physiol. 2019;10:361.

23. Forte N, Imperatore R, Marfella B, Nicois A, Verde R, Palomba L, et al. The Blue Light-Responsive Lateral Pathway of the Retinohypothalamic Tract Promotes Endocannabinoid-Driven Modulation of Orexin Neurons. J Neurochem. 2025;169(6):e70137.

24. Di Marzo V. New approaches and challenges to targeting the endocannabinoid system. Nat Rev Drug Discov. 2018;17(9):623–39.

25. Niepokny TD, Frey-Burkart H, Mintz EM. Temporal and spatial layout of endocannabinoid system components in the mouse suprachiasmatic nucleus. Neuroscience. 2025;564:179–93.

26. Sladek M, Houdek P, Sumova A. Circadian profiling reveals distinct regulation of endocannabinoid system in the rat plasma, liver and adrenal glands by light-dark and feeding cycles. Biochim Biophys Acta Mol Cell Biol Lipids. 2019;1864(12):158533.

27. Farfara D, Lewitus GM, Korin B, Hajjo H, Feder CL, Sulimani L, et al. Circadian-related Dynamics of the Endocannabinoid System in Male Mouse Brain. bioRxiv. 2025:2025.04.15.648929.

28. Lefort C, Roumain M, Van Hul M, Rastelli M, Manco R, Leclercq I, et al. Hepatic NAPE-PLD Is a Key Regulator of Liver Lipid Metabolism. Cells. 2020;9(5).

29. Zarrow JE, Alli-Oluwafuyi AM, Youwakim CM, Kim K, Jenkins AN, Suero IC, et al. Small Molecule Activation of NAPE-PLD Enhances Efferocytosis by Macrophages. ACS Chem Biol. 2023;18(8):1891–904.

30. Fischer C, Thomas D, Gurke R, Tegeder I. Brain region specific regulation of anandamide (down) and sphingosine-1-phosphate (up) in association with anxiety (AEA) and resilience (S1P) in a mouse model of chronic unpredictable mild stress. Pflugers Arch. 2024;476(12):1863–80.

31. Karanian DA, Karim SL, Wood JT, Williams JS, Lin S, Makriyannis A, et al. Endocannabinoid enhancement protects against kainic acid-induced seizures and associated brain damage. J Pharmacol Exp Ther. 2007;322(3):1059–66.

32. Shubina L, Aliev R, Kitchigina V. Endocannabinoid-dependent protection against kainic acid-induced long-term alteration of brain oscillations in guinea pigs. Brain Res. 2017;1661:1–14.

33. Natarajan V, Schmid PC, Schmid HH. N-acylethanolamine phospholipid metabolism in normal and ischemic rat brain. Biochim Biophys Acta. 1986;878(1):32–41.

34. Fornelos N, Franzosa EA, Bishai J, Annand JW, Oka A, Lloyd-Price J, et al. Growth effects of N-acylethanolamines on gut bacteria reflect altered bacterial abundances in inflammatory bowel disease. Nat Microbiol. 2020;5(3):486–97.

35. Berdyshev EV, Schmid PC, Dong Z, Schmid HH. Stress-induced generation of N-acylethanolamines in mouse epidermal JB6 P+ cells. Biochem J. 2000;346 Pt 2(Pt 2):369–74.

36. Mallipedhi A, Prior SL, Dunseath G, Bracken RM, Barry J, Caplin S, et al. Changes in plasma levels of N-arachidonoyl ethanolamine and N-palmitoylethanolamine following bariatric surgery in morbidly obese females with impaired glucose homeostasis. J Diabetes Res. 2015;2015:680867.

37. Veilleux A, Di Marzo V, Silvestri C. The Expanded Endocannabinoid System/Endocannabinoidome as a Potential Target for Treating Diabetes Mellitus. Curr Diab Rep. 2019;19(11):117.

38. Keppel Hesselink JM, de Boer T, Witkamp RF. Palmitoylethanolamide: A Natural Body-Own Anti-Inflammatory Agent, Effective and Safe against Influenza and Common Cold. Int J Inflam. 2013;2013:151028.

39. Epps DE, Schmid PC, Natarajan V, Schmid HH. N-Acylethanolamine accumulation in infarcted myocardium. Biochem Biophys Res Commun. 1979;90(2):628–33.

40. Ponomarenko AI, Tyrtyshnaia AA, Pislyagin EA, Dyuizen IV, Sultanov RM, Manzhulo IV. N-docosahexaenoylethanolamine reduces neuroinflammation and cognitive impairment after mild traumatic brain injury in rats. Sci Rep. 2021;11(1):756.

41. Chen R, Zuo Z, Li Q, Wang H, Li N, Zhang H, et al. DHA substitution overcomes high-fat diet-induced disturbance in the circadian rhythm of lipid metabolism. Food Funct. 2020;11(4):3621–31.

42. Igarashi M, DiPatrizio NV, Narayanaswami V, Piomelli D. Feeding-induced oleoylethanolamide mobilization is disrupted in the gut of diet-induced obese rodents. Biochim Biophys Acta. 2015;1851(9):1218–26.

43. Pacher P, Kunos G. Modulating the endocannabinoid system in human health and disease--successes and failures. FEBS J. 2013;280(9):1918–43.

44. Pacher P, Batkai S, Kunos G. The endocannabinoid system as an emerging target of pharmacotherapy. Pharmacol Rev. 2006;58(3):389–462.

45. Engeli S. Dysregulation of the endocannabinoid system in obesity. J Neuroendocrinol. 2008;20 Suppl 1:110–5.

46. Castonguay-Paradis S, Lacroix S, Rochefort G, Parent L, Perron J, Martin C, et al. Dietary fatty acid intake and gut microbiota determine circulating endocannabinoidome signaling beyond the effect of body fat. Sci Rep. 2020;10(1):15975.

47. Castonguay-Paradis S, Parent L, St-Arnaud G, Perron J, Dumais E, Flamand N, et al. The Human Fecal Endocannabinoidome Mediator Profile Is Mainly Defined by the Fecal Microbiota and Diet. J Clin Endocrinol Metab. 2025;110(3):739–47.

48. Roussel C, Lessard-Lord J, Nallabelli N, Muller C, Flamand N, Silvestri C, et al. Human Gut Microbes Produce EPA- and DHA-Derived Oxylipins, but not N-Acyl-Ethanolamines, From Fish Oil. FASEB J. 2025;39(12):e70713.

49. Manca C, Boubertakh B, Leblanc N, Deschenes T, Lacroix S, Martin C, et al. Germ-free mice exhibit profound gut microbiota-dependent alterations of intestinal endocannabinoidome signaling. J Lipid Res. 2020;61(1):70–85.

50. Thaiss CA, Zeevi D, Levy M, Zilberman-Schapira G, Suez J, Tengeler AC, et al. Transkingdom control of microbiota diurnal oscillations promotes metabolic homeostasis. Cell. 2014;159(3):514–29.

51. Bloemen JG, Venema K, van de Poll MC, Olde Damink SW, Buurman WA, Dejong CH. Short chain fatty acids exchange across the gut and liver in humans measured at surgery. Clin Nutr. 2009;28(6):657–61.

52. Matenchuk BA, Mandhane PJ, Kozyrskyj AL. Sleep, circadian rhythm, and gut microbiota. Sleep Med Rev. 2020;53:101340.

53. Callahan BJ, McMurdie PJ, Rosen MJ, Han AW, Johnson AJ, Holmes SP. DADA2: High-resolution sample inference from Illumina amplicon data. Nat Methods. 2016;13(7):581–3.

54. Yilmaz P, Parfrey LW, Yarza P, Gerken J, Pruesse E, Quast C, et al. The SILVA and “All-species Living Tree Project (LTP)” taxonomic frameworks. Nucleic Acids Res. 2014;42(Database issue):D643–8.

55. Lu Y, Zhou G, Ewald J, Pang Z, Shiri T, Xia J. MicrobiomeAnalyst 2.0: comprehensive statistical, functional and integrative analysis of microbiome data. Nucleic Acids Res. 2023;51(W1):W310–W8.

56. Bligh EG, Dyer WJ. A rapid method of total lipid extraction and purification. Can J Biochem Physiol. 1959;37(8):911–7.

57. Turcotte C, Archambault AS, Dumais E, Martin C, Blanchet MR, Bissonnette E, et al. Endocannabinoid hydrolysis inhibition unmasks that unsaturated fatty acids induce a robust biosynthesis of 2-arachidonoyl-glycerol and its congeners in human myeloid leukocytes. FASEB J. 2020;34(3):4253–65.

58. Everard A, Plovier H, Rastelli M, Van Hul M, de Wouters d’Oplinter A, Geurts L, et al. Intestinal epithelial N-acylphosphatidylethanolamine phospholipase D links dietary fat to metabolic adaptations in obesity and steatosis. Nat Commun. 2019;10(1):457.

59. Schebb NH, Kampschulte N, Hagn G, Plitzko K, Meckelmann SW, Ghosh S, et al. Technical recommendations for analyzing oxylipins by liquid chromatography-mass spectrometry. Sci Signal. 2025;18(887):eadw1245.

60. Foster ZS, Sharpton TJ, Grunwald NJ. Metacoder: An R package for visualization and manipulation of community taxonomic diversity data. PLoS Comput Biol. 2017;13(2):e1005404.

61. Parkar SG, Kalsbeek A, Cheeseman JF. Potential Role for the Gut Microbiota in Modulating Host Circadian Rhythms and Metabolic Health. Microorganisms. 2019;7(2).

62. Sun Y, Zeng X, Liu Y, Zhan S, Wu Z, Zheng X, et al. Dendrobium officinale polysaccharide attenuates cognitive impairment in circadian rhythm disruption mice model by modulating gut microbiota. Int J Biol Macromol. 2022;217:677–88.

63. Kim YM, Snijders AM, Brislawn CJ, Stratton KG, Zink EM, Fansler SJ, et al. Light-Stress Influences the Composition of the Murine Gut Microbiome, Memory Function, and Plasma Metabolome. Front Mol Biosci. 2019;6:108.

64. Heddes M, Altaha B, Niu Y, Reitmeier S, Kleigrewe K, Haller D, et al. The intestinal clock drives the microbiome to maintain gastrointestinal homeostasis. Nat Commun. 2022;13(1):6068.

65. Parker BJ, Wearsch PA, Veloo ACM, Rodriguez-Palacios A. The Genus Alistipes: Gut Bacteria With Emerging Implications to Inflammation, Cancer, and Mental Health. Front Immunol. 2020;11:906.

66. Lin TC, Soorneedi A, Guan Y, Tang Y, Shi E, Moore MD, et al. Turicibacter fermentation enhances the inhibitory effects of Antrodia camphorata supplementation on tumorigenic serotonin and Wnt pathways and promotes ROS-mediated apoptosis of Caco-2 cells. Front Pharmacol. 2023;14:1203087.

67. Xi L, Wen X, Jia T, Han J, Qin X, Zhang Y, et al. Comparative study of the gut microbiota in three captive Rhinopithecus species. BMC Genomics. 2023;24(1):398.

68. Yu Z, Li D, Sun H. Herba Origani alleviated DSS-induced ulcerative colitis in mice through remolding gut microbiota to regulate bile acid and short-chain fatty acid metabolisms. Biomed Pharmacother. 2023;161:114409.

69. Chen Y, Ye L, Zhu J, Chen L, Chen H, Sun Y, et al. Disrupted Tuzzerella abundance and impaired L-glutamine levels induce Treg accumulation in ovarian endometriosis: a comprehensive multi-omics analysis. Metabolomics. 2024;20(2):32.

70. Zhou D, Pan Q, Xin FZ, Zhang RN, He CX, Chen GY, et al. Sodium butyrate attenuates high-fat diet-induced steatohepatitis in mice by improving gut microbiota and gastrointestinal barrier. World J Gastroenterol. 2017;23(1):60–75.

71. Shimizu Y, Yamamura R, Yokoi Y, Ayabe T, Ukawa S, Nakamura K, et al. Shorter sleep time relates to lower human defensin 5 secretion and compositional disturbance of the intestinal microbiota accompanied by decreased short-chain fatty acid production. Gut Microbes. 2023;15(1):2190306.

72. Kim CH. Complex regulatory effects of gut microbial short-chain fatty acids on immune tolerance and autoimmunity. Cell Mol Immunol. 2023;20(4):341–50.

73. Suriano F, Manca C, Flamand N, Van Hul M, Delzenne NM, Silvestri C, et al. A Lipidomics- and Transcriptomics-Based Analysis of the Intestine of Genetically Obese (ob/ob) and Diabetic (db/db) Mice: Links with Inflammation and Gut Microbiota. Cells. 2023;12(3).

74. Froy O, Weintraub Y. The circadian clock, metabolism, and inflammation-the holy trinity of inflammatory bowel diseases. Clin Sci (Lond). 2025;139(13):777–90.

75. Salaun C, Courvalet M, Rousseau L, Cailleux K, Breton J, Bole-Feysot C, et al. Sex-dependent circadian alterations of both central and peripheral clock genes expression and gut-microbiota composition during activity-based anorexia in mice. Biol Sex Differ. 2024;15(1):6.

76. Ma T, Matsuo R, Kurogi K, Miyamoto S, Morita T, Shinozuka M, et al. Sex-dependent effects of chronic jet lag on circadian rhythm and metabolism in mice. Biol Sex Differ. 2024;15(1):102.

77. Di Minno MN, Russolillo A, Lupoli R, Ambrosino P, Di Minno A, Tarantino G. Omega-3 fatty acids for the treatment of non-alcoholic fatty liver disease. World J Gastroenterol. 2012;18(41):5839–47.

78. Kotsos D, Tziomalos K. Microsomal Prostaglandin E Synthase-1 and -2: Emerging Targets in Non-Alcoholic Fatty Liver Disease. Int J Mol Sci. 2023;24(3).

79. Loomba R, Morgan E, Yousefi K, Li D, Geary R, Bhanot S, et al. Antisense oligonucleotide DGAT-2 inhibitor, ION224, for metabolic dysfunction-associated steatohepatitis (ION224-CS2): results of a 51-week, multicentre, randomised, double-blind, placebo-controlled, phase 2 trial. Lancet. 2025;406(10505):821–31.

80. Yadav A, Ouyang X, Barkley M, Watson JC, Madamanchi K, Kramer J, et al. Regulation of lipid dysmetabolism and neuroinflammation progression linked with Alzheimer’s disease through modulation of Dgat2. bioRxiv. 2025:2025.02.18.638929.

81. McFie PJ, Patel A, Stone SJ. The monoacylglycerol acyltransferase pathway contributes to triacylglycerol synthesis in HepG2 cells. Sci Rep. 2022;12(1):4943.

82. Ben Necib R, Manca C, Lacroix S, Martin C, Flamand N, Di Marzo V, et al. Hemp seed significantly modulates the endocannabinoidome and produces beneficial metabolic effects with improved intestinal barrier function and decreased inflammation in mice under a high-fat, high-sucrose diet as compared with linseed. Frontiers in Immunology. 2022;Volume 13 - 2022.

83. Lee JW, Huang BX, Kwon H, Rashid MA, Kharebava G, Desai A, et al. Orphan GPR110 (ADGRF1) targeted by N-docosahexaenoylethanolamine in development of neurons and cognitive function. Nat Commun. 2016;7:13123.

84. Lambert DM, Vandevoorde S, Diependaele G, Govaerts SJ, Robert AR. Anticonvulsant activity of N-palmitoylethanolamide, a putative endocannabinoid, in mice. Epilepsia. 2001;42(3):321–7.

85. Suardiaz M, Estivill-Torrus G, Goicoechea C, Bilbao A, Rodriguez de Fonseca F. Analgesic properties of oleoylethanolamide (OEA) in visceral and inflammatory pain. Pain. 2007;133(1-3):99–110.

86. Kathuria S, Gaetani S, Fegley D, Valino F, Duranti A, Tontini A, et al. Modulation of anxiety through blockade of anandamide hydrolysis. Nat Med. 2003;9(1):76–81.

87. Karatas H, Cakir-Aktas C. 12/15 Lipoxygenase as a Therapeutic Target in Brain Disorders. Noro Psikiyatr Ars. 2019;56(4):288–91.

88. Niepokny TD, Mintz EM. A Cannabinoid Receptor 1 Agonist Reduces Light-induced Phase Delays in Male But Not Female Mice. J Biol Rhythms. 2023;38(4):358–65.

89. Herrera MI, Kolliker-Frers R, Barreto G, Blanco E, Capani F. Glial Modulation by N-acylethanolamides in Brain Injury and Neurodegeneration. Front Aging Neurosci. 2016;8:81.

90. Wasilewski M, Wieckowski MR, Dymkowska D, Wojtczak L. Effects of N-acylethanolamines on mitochondrial energetics and permeability transition. Biochim Biophys Acta. 2004;1657(2-3):151–63.

91. Movsesyan VA, Stoica BA, Yakovlev AG, Knoblach SM, Lea PMt, Cernak I, et al. Anandamide-induced cell death in primary neuronal cultures: role of calpain and caspase pathways. Cell Death Differ. 2004;11(10):1121–32.

92. Giuffrida A, Leweke FM, Gerth CW, Schreiber D, Koethe D, Faulhaber J, et al. Cerebrospinal anandamide levels are elevated in acute schizophrenia and are inversely correlated with psychotic symptoms. Neuropsychopharmacology. 2004;29(11):2108–14.

93. Ashton A, Jagannath A. Disrupted Sleep and Circadian Rhythms in Schizophrenia and Their Interaction With Dopamine Signaling. Front Neurosci. 2020;14:636.

94. Van Drunen R, Eckel-Mahan K. Circadian rhythms as modulators of brain health during development and throughout aging. Front Neural Circuits. 2022;16:1059229.

95. Salahuddin MF, Bugingo R, Mahdi F, Spencer D, Manzar MD, Paris JJ. Physiological and Psychological Impacts of Shift Work Among Student Pharmacists: Sex Differences in Stress and Health Outcomes. Psychiatry International. 2025;6(2):47.

96. Morrison M, Halson SL, Weakley J, Hawley JA. Sleep, circadian biology and skeletal muscle interactions: Implications for metabolic health. Sleep Med Rev. 2022;66:101700.

97. Mansingh S, Maier G, Delezie J, Westermark PO, Ritz D, Duchemin W, et al. More than the clock: distinct regulation of muscle function and metabolism by PER2 and RORalpha. J Physiol. 2024;602(23):6373–402.

98. McCarthy JJ, Andrews JL, McDearmon EL, Campbell KS, Barber BK, Miller BH, et al. Identification of the circadian transcriptome in adult mouse skeletal muscle. Physiol Genomics. 2007;31(1):86–95.

99. Harfmann BD, Schroder EA, Esser KA. Circadian rhythms, the molecular clock, and skeletal muscle. J Biol Rhythms. 2015;30(2):84–94.

100. Riley DA, Bain JL, Thompson JL, Fitts RH, Widrick JJ, Trappe SW, et al. Decreased thin filament density and length in human atrophic soleus muscle fibers after spaceflight. J Appl Physiol (1985). 2000;88(2):567–72.

101. Lu L, Williams G, Doherty P. 2-Linoleoylglycerol Is a Partial Agonist of the Human Cannabinoid Type 1 Receptor that Can Suppress 2-Arachidonolyglycerol and Anandamide Activity. Cannabis and Cannabinoid Research. 2019;4(4):231–9.

102. Osei-Hyiaman D, DePetrillo M, Pacher P, Liu J, Radaeva S, Batkai S, et al. Endocannabinoid activation at hepatic CB1 receptors stimulates fatty acid synthesis and contributes to diet-induced obesity. J Clin Invest. 2005;115(5):1298–305.

103. Tedesco L, Valerio A, Cervino C, Cardile A, Pagano C, Vettor R, et al. Cannabinoid type 1 receptor blockade promotes mitochondrial biogenesis through endothelial nitric oxide synthase expression in white adipocytes. Diabetes. 2008;57(8):2028–36.

104. Serhan CN. Pro-resolving lipid mediators are leads for resolution physiology. Nature. 2014;510(7503):92–101.

105. Calder PC. n-3 polyunsaturated fatty acids, inflammation, and inflammatory diseases. Am J Clin Nutr. 2006;83(6 Suppl):1505S–19S.

106. Vegiopoulos A, Muller-Decker K, Strzoda D, Schmitt I, Chichelnitskiy E, Ostertag A, et al. Cyclooxygenase-2 controls energy homeostasis in mice by de novo recruitment of brown adipocytes. Science. 2010;328(5982):1158–61.

107. Edwards M, Mohiuddin SS. Biochemistry, Lipolysis. StatPearls. Treasure Island (FL) ineligible companies. Disclosure: Shamim Mohiuddin declares no relevant financial relationships with ineligible companies.2025.

108. Dyar KA, Lutter D, Artati A, Ceglia NJ, Liu Y, Armenta D, et al. Atlas of Circadian Metabolism Reveals System-wide Coordination and Communication between Clocks. Cell. 2018;174(6):1571–85 e11.

109. Alvarez JD, Hansen A, Ord T, Bebas P, Chappell PE, Giebultowicz JM, et al. The circadian clock protein BMAL1 is necessary for fertility and proper testosterone production in mice. J Biol Rhythms. 2008;23(1):26–36.

110. Sellix MT. Clocks underneath: the role of peripheral clocks in the timing of female reproductive physiology. Front Endocrinol (Lausanne). 2013;4:91.

111. Mohammadpour-Asl S, Roshan-Milani S, Fard AA, Golchin A. Hormetic effects of a cannabinoid system component, 2-arachidonoyl glycerol, on cell viability and expression profile of growth factors in cultured mouse Sertoli cells: Friend or foe of male fertility? Reprod Toxicol. 2024;125:108575.

112. Sanchez AM, Cioffi R, Vigano P, Candiani M, Verde R, Piscitelli F, et al. Elevated Systemic Levels of Endocannabinoids and Related Mediators Across the Menstrual Cycle in Women With Endometriosis. Reprod Sci. 2016;23(8):1071–9.

113. Tamura H, Takasaki A, Miwa I, Taniguchi K, Maekawa R, Asada H, et al. Oxidative stress impairs oocyte quality and melatonin protects oocytes from free radical damage and improves fertilization rate. J Pineal Res. 2008;44(3):280–7.

114. Sciarra F, Franceschini E, Campolo F, Gianfrilli D, Pallotti F, Paoli D, et al. Disruption of Circadian Rhythms: A Crucial Factor in the Etiology of Infertility. Int J Mol Sci. 2020;21(11).

115. Abo SMC, Layton AT. Modeling the circadian regulation of the immune system: Sexually dimorphic effects of shift work. PLoS Comput Biol. 2021;17(3):e1008514.

116. Arjona A, Sarkar DK. The circadian gene mPer2 regulates the daily rhythm of IFN-gamma. J Interferon Cytokine Res. 2006;26(9):645–9.

117. Cayetanot F, Nygard M, Perret M, Kristensson K, Aujard F. Plasma levels of interferon-gamma correlate with age-related disturbances of circadian rhythms and survival in a non-human primate. Chronobiol Int. 2009;26(8):1587–601.

